# Replisome progression regulates R-loop mediated transcriptional repression

**DOI:** 10.1101/2025.07.29.667376

**Authors:** Ioannis Tsirkas, Clare S.K. Lee, Daniel Dovrat, Neha Singh, Zohar Paleiov, Maxime Lalonde, Atiqa Sajid, Amir Aharoni, Stephan Hamperl

**Affiliations:** Chromosome Dynamics and Genome Stability, Institute of Epigenetics and Stem Cells, Helmholtz Munich, Munich 81377, Germany; Department of Life Sciences, Ben-Gurion University of the Negev, Be’er Sheva 84105, Israel

## Abstract

Maintaining cellular proliferation necessitates the synchronized activity of diverse molecular machineries operating in parallel on the genome. Wide-spread transcription and R-loop formation can interfere with genome duplication, causing transcription-replication conflicts (TRCs). Here, we use live-cell imaging for simultaneous monitoring of replication fork progression and transcription dynamics of an R-loop prone gene. While robust replisome progression through R-loops is observed in wild-type cells, it is impaired in RnaseH, Mph1 and Sen1 mutants. Intriguingly, we find that R-loop formation inhibits gene transcription, but this inhibition is reversed by replisome passage, demonstrating the dynamic crosstalk between replication, transcription and R-loops in a single cell cycle. Unexpectedly, we find that R-loops also have beneficial roles in reducing the density of RNAPII molecules on the gene, thereby preventing fork stalling upon high RNAPII occupancy. These findings illuminate that regulating R-loops and RNAPII density together is critical for preventing harmful TRCs.

## INTRODUCTION

R-loops are non-B DNA structures formed co-transcriptionally when the newly-synthesized RNA strand anneals with its complementary DNA strand, displacing the single-stranded coding DNA [1]. Although R-loops maintain a variety of physiological functions, they also constitute DNA damage hotspots, contributing to genomic instability and mutagenesis [2–4]. Recent discoveries have connected R-loop persistence with several human diseases such as oncogenesis and neurological disorders, establishing their clinical significance as diagnostic tool and potential biomarkers [5,6]. R-loop mapping studies have revealed hundreds to thousands of R-loop prone genomic sequences [7–9]. Considering R-loop abundance and the challenge they pose for genomic integrity, cells need to tightly coordinate their formation with other essential processes such as DNA replication.

DNA replication and transcription machineries ensure faithful duplication of the genome and proper gene expression, respectively. However, both types of machinery share the same DNA template. During S-phase, a replisome may confront multiple RNA polymerases (RNAP) occupying highly transcribed genes, leading to Transcription-Replication Conflicts (TRCs) [10–12]. Although the two processes are generally segregated both spatially and temporally [13], it has been shown that TRCs can occur frequently in eukaryotic cells [14,15]. If not resolved, they can result in replication fork stalling or collapse, causing genomic instability and increased frequency of DNA double-strand breaks (DSB) [16–18]. TRCs are imminent threats to genomic integrity, especially in the event where the transcribing RNAP is facing the incoming replisome head-on (HO) [19,20]. Co-directional (CoD) conflicts are better tolerated by the cells, but can also show detrimental outcomes, particularly when replication forks are challenged with more persistent transcriptional blocks such as R-loops [20]. Thus, understanding how cells avoid and overcome such hazardous R-loop-dependent TRCs is of paramount importance for deciphering the pathways that contribute to genome integrity.

Apart from their impact on replisome progression, R-loops were shown *in vitro* and *in vivo* to impede transcription by interfering with RNAPII progression, resulting in early transcription termination [21–23]. Arrest of RNAPII’s progression by R-loops can eventually lead to their eviction and sequential degradation [24,25]. Furthermore, thousands of genes have been identified with skewed GC content at their 3’ ends [26], consistent with a role of R-loops in transcription termination. Thus, R-loops can have both a negative impact on genomic stability and, on the other hand, regulatory roles during genome duplication and transcription.

Considering the disruptive potential of R-loops, cells have evolved an extensive plethora of protein factors to counteract RNA:DNA hybrid accumulation. This includes proteins that enzymatically degrade and remove RNA:DNA hybrids from the genome, such as the family of Ribonuclease H (RNH) enzymes [27] or RNA:DNA helicases such as Sen1 or Mph1 designated to unwind R-loops and maintain unhindered replisome progression and gene transcription [28] . This redundancy suggests that R-loop levels need to be tightly regulated to preserve proper gene expression, replisome progression and cellular function [29]. One main limitation in our understanding of the interplay between R-loops, transcription and replication is the lack of techniques for the simultaneous measurement of fork progression and transcription dynamics at an inducible R-loop gene in a single cell cycle.

Here, we took advantage of a previously established system for the real-time monitoring of transcription and replication at an individual locus in single live cells [30]. By inserting the inducible, highly R-loop prone mouse Antisense of IGF2R non-protein coding RNA (mAIRN) sequence in the path of a single replisome, we found that R-loop formation strongly suppresses mAIRN transcription. Interestingly, replisome progression through mAIRN leads to a strong recovery of transcriptional activity. This observation suggests a replisome-dependent regulation of transcriptional activity at R-loop forming genes. We further found that in wildtype (WT) cells, replisome progression is robust but is slowed down in RNH, Mph1 and Sen1 mutant cells. Unexpectedly, we provide evidence that R-loops can also have a beneficial role to reduce RNAPII occupancy on highly expressed genes to allow normal replication fork progression. Overall, our findings offer new insights into the functional interplay between R-loop formation, transcriptional regulation, and replisome, and reveal how cells coordinate these processes to ensure the suppression of harmful TRCs.

## MATERIAL AND METHODS

### Reagents and biological materials

A detailed list of yeast and bacterial strains (**Table S1**), plasmids (**Table S2**), oligonucleotides (**Table S3**), reagents, commercial kits and assays (**Table S4**), and software and algorithms (**Table S5**) used in this study can be found in the Supplemental Information.

### Saccharomyces cerevisiae strains generation

All yeast strains used in this study were generated on the background of W1588, which is identical to W303 but with a wild-type copy of Rad5, MATa *S. cerevisiae* strain, integrated with *lacO*x256 and *tetO*x224 arrays at *chrIV*:332960 and *chrIV*:352560, respectively, adjacent to *ARS413.* LacI-HaloTag and tetR-tdTomato fusion proteins are expressed in the nucleus to visualize the arrays as distinct fluorescent foci[30]. The mAIRN sequence (1401 bp) was expressed from the unidirectional GAL10 promoter (Pgal, 404 bp) and labeled with 14xPP7. Then, this mAIRN reporter cassette was flanked by CUT60 and ADH1 terminator sequences and the whole cassette was amplified and inserted between the two arrays at *chrIV*:336186 in either CoD or HO orientation relative to replisome emerging from *ARS413*. The mAIRN reporter cassette integration was performed using a marker-free CRISPR-Cas9 approach, targeting *natMX* antibiotic cassettes with specific gRNAs [31].

For Rnh1 overexpression, RNH1orf was expressed by the Pgal (454 bp) and 3 copies of HAtag were fused to its C’-terminus, followed by CYC1 terminator, in a URA3-integrative plasmid. The generation of *rnh1rnh201*ΔΔ and *mph1*Δ yeast strains was performed by replacing the respective genes with *natMX*, *hphMX*, or *kanMX* antibiotic cassettes. The *sen1-3* mutant (W773A, E774A, W777A) strains [32] were generated with CRISPR-Cas9, using a gRNA (gRNA sequence: TAAAATTCTGGGAATCATGT) which targets the SEN1 gene in the mutated region, using a DNA donor containing the mutation with 80 bp homology to the genome on each side of the targeted region. The *rpb1-1* mutant (G4622A) strains [33] were generated with CRISPR-Cas9, as described before [30]. MNase fusion of RPB2 was performed by amplification of the MNase-3xHA-KanMX cassette from a plasmid and integration in the C’-end of RPB2 gene, excluding the native termination codon. All integrations, replacements and mutations were validated by PCR and Sanger sequencing.

### Microscopy

Yeast cells were grown overnight in synthetic complete (SC) Medium containing 2% raffinose (RAF) at 30°C. Yeast cultures were diluted at OD_600_ = 0.2 and SiR-HALO dye was added to a final concentration of 800 nM. One hour following SiR-HALO dye addition, 10 μg/ml α-factor was added to arrest the cells in G1 phase, and the cultures were incubated for two additional hours. Cells were then immobilized on microscopy slide chambers (Ibidi) coated with 2 mg/ml concanavalin A for 10-15 min and washed thoroughly from α-factor and SiR-HALO dye with warm SC medium containing 2% RAF prior to microscopy experiments. For the induction of mAIRN transcription, 2% galactose (GAL) was added to the culture media 2h before and during the microscope experiment. Live-cell imaging of the cells was performed on an Axio Observer inverted wide-field microscope (Zeiss) with a Colibri 7 LED light source, at 1 min intervals for 4 hr at 30 °C, using a x63 oil objective (NA = 1.4) in 3D (8 z-sections, 0.8 μm apart). LacI-Halo-SiR and tetR-tdTomato were excited with 650 and 561 nm illumination, respectively. PCP-Envy was excited with 488 nm illumination.

### Western blot analysis

For the detection of Rnh1-3xHA or Rpb2-MNase-3xHA expression by western blot, 5 ml of yeast cells were grown until OD_600_ ∼ 0.5) in 2% RAF. For induction of PP7-mAIRN and Rnh1 OE, 2% GAL was added in in the media and cells were grown for an additional 2 hr (OD_600_ ∼ 1). 2-3 ml of cell cultures were transferred into separate 1.5 ml tubes and centrifuged at >20,000 xg for 1 min. The supernatants were discarded and cell pellets were snap-frozen in liquid N_2_ and stored in - 80 °C overnight.

The next day, the pellets were resuspended in 500 μl ice-cold ddH_2_O and 75 μl NaOH/β-ME solution (1.85 M NaOH and 7.5% β-mercaptoethanol). The cell solutions were vortexed for 4 s and incubated on ice for 15 min. Then, 75 μl 55% TCA was also added, the tubes were vortexed for 4 s and incubated on ice for 10 min. Lysed cells were centrifuged for 10 min at >20,000 xg and at 4 °C and the supernatant was discarded. Finally, tubes were centrifuged for another 30 s for 10 min at >20,000 xg at 4 °C to collect and discard completely the residual liquid. Pellets were resuspended with 30 - 40 μl of High Urea (HU) protein loading buffer (8 M urea, 5% SDS, 200 mM Tris-HCl pH = 6.8, 1 mM EDTA, 0.1% bromophenol blue). If the suspensions turned green or yellow colour, 1 μl 1 M Tris-HCl pH = 8 was added. Proteins were denatured at 65 °C for 10 min with mild shaking.

4-10 μl of protein lysates were loaded and run on a 10% SDS-PAGE gel for 1.5 - 2 hr at 120 V. Separated protein bands were transferred to a PVDF membrane followed by blocking with PBST + 10% skim milk overnight. The membrane was washed with PBST once for 5 min, and was incubated with a rat anti-HA antibody conjugated with HRP enzyme (1:2000) in PBST+10% skim milk for 1 hr. The membrane was washed three times with PBST and HA-labelled Rnh1 or Rpb2 were detected with the EZ-ECL Chemiluminescence detection kit (Thermo Fisher Scientific) according to the manufacturer’s protocol. Detection of RNAPII bands was performed with a primary rabbit anti-RNAPII antibody (1:2000) and a secondary goat-anti-rabbit antibody (1:10000) conjugated with HRP.

### Protein crosslinking

Yeast cells were grown overnight in 2% RAF (Yeast Peptone Raffinose -YPR) and then diluted at 50 ml and OD_600_ ∼ 0.1 - 0.2 with prewarmed YPR in 100 ml Erlenmeyer flasks. Cell cultures were grown until OD_600_ ∼ 0.5, then 2% GAL was added in the media and the cultures were grown for another 2 hr, until OD_600_ ∼ 1. For experiments with cell cycle synchronization, overnight cell cultures were diluted at 100 ml and OD_600_ ∼ 0.1 - 0.2 in 250 ml flasks and they were grown until OD_600_ ∼ 0.5 or 0.6. Then, 10 μg/ml α-factor was added together with 2% GAL and cells were incubated for another 2 hr, until OD_600_ ∼ 1. Then, cells were split into 50 ml tubes and were centrifuged in a 30 °C prewarmed centrifuge for 4 min at 1000 xg. The supernatants were discarded and cells of the same strain were collected in a single 50 ml tube. Cells were washed from α-factor twice with 45 ml prewarmed YPR + 2% GAL, and they were resuspended in 100 ml warm YPR + 2% GAL. 45 ml of yeast cultures were sampled at desired timepoints (10 min and 50 min post release into S phase).

For crosslinking, 3 ml of 16% formaldehyde (FA) was added dropwise in 45 ml of yeast cultures (final concentration 1%) and cells were fixed in 100 ml flasks for 15 minutes at 30 °C with shaking. FA was quenched with 2.6 ml of 2.5 M glycine (final concentration 128 mM) for 5 min minutes at 30°C. Cells were transferred and harvested in 50 ml tubes (1000 ×g, 2 min at 4 °C), and washed once with ice-cold PBS. After this step, fixed cells were always treated on ice. Washed cells were centrifuged (1000 × g, 2 min at 4 °C) and cell pellets were resuspended in 1 ml of ddH_2_O and transferred into 1.5 ml tubes. Yeast cells were collected by centrifugation at 20000 xg for 1:30 min at 4 °C in a microcentrifuge, the supernatants were discarded and the pellets were frozen in liquid N_2_ and stored at -80 °C for at least overnight.

### RNAPII Chromatin Immunoprecipitation (ChIP)-qPCR

ChIP-qPCR experiments were performed as described before[34]. Samples and buffers were kept on ice at all times. Briefly, frozen fixed cell pellets were washed with 500 µl Lysis buffer [50 mM HEPES pH = 7.5, 140 mM NaCl, 5 mM EDTA pH = 8, 5mM EGTA pH = 8, 1% Triton X-100 (w/v), 0.1% DOC (w/v), 1x protease and phosphatase inhibitors] and then they were centrifuged at 20000 xg for 1:30 min at 4°C. The supernatants were discarded and cell pellets were resuspended with 500 µl Lysis buffer. ∼750 μl precooled glass beads (1 mm) were added to cover the whole suspension and the tubes were filled with Lysis Buffer. Cells were disrupted on a VXR basic IKA Vibrax orbital shaker at 2200 rpm at 4 °C for 15 min, three times, with 10 min breaks on ice in between. The bottom and top of the 1.5 mL tube were pierced using a hot needle and placed into a 15 mL tube. After centrifugation (130 xg, 2 min at 4 °C), the lysates were collected in the 15 mL tubes while the glass beads stayed in the tube. The volume of the suspension was increased to 1 mL with Lysis buffer and transferred to a 1 mL Covaris sonication glass tube. Sonification was performed on a Covaris instrument (25 min, Peak Incident Power: 140 W, Duty Factor: 5%, Cycles/Burst: 200). Afterwards, the sheared chromatin was cleared by centrifugation (20 min, 20000 xg at 4 ^ο^C) and the supernatant was transferred to a DNA-low-binding 1.5 mL tube. The solution was split into two aliquots: a total of 90 μL served as input (IN) control, and 810 μL was used for IP.

40 μl protein A magnetic beads were washed thrice with Lysis Buffer at 4 °C and were resuspended with the 810 μl of sheared chromatin. The solution was filled up to 990 μl with Lysis Buffer, supplemented with 5 μg (10 μl) anti-RNAPII CTD antibody (8WG15 clone) and was incubated for 2 hr at 4 °C on a rotating platform. After IP, the beads were washed three times with lysis buffer, twice with Wash buffer I [50 mM HEPES pH = 7.5, 500 mM NaCl, 2 mM EDTA, 1% (w/v) Triton X-100, 0.1% (w/v) DOC], twice with Wash buffer II [10 mM Tris-HCl pH = 8.0, 250 mM LiCl, 2 mM EDTA, 0.5 % (v/v) Igepal CA-630, 0.5% (w/v) DOC], and once with TE buffer (10 mM Tris-HCl pH = 8.0, 1 mM EDTA). A total of 390 μL or 300 μL of IRN Buffer (50 mM Tris–HCl pH = 8.0, 20 mM EDTA, 500 mM NaCl) were added to the IP beads or the IN samples, respectively. IP and IN samples were incubated with 10 μL RNAse A (10 mg/mL) at 37 °C for 30 min. Then, SDS was added to a final concentration of 0.5% together with 10 μL Proteinase K (10 mg/mL) and the samples were incubated for 1 hr at 56 °C. Crosslinking was reversed at 65 °C overnight. The next day, 1 ng spike-in plasmids (K18) were added in each tube and DNA was isolated by phenol-chloroform extraction. Samples were washed with 70% EtOH, were resuspensed in 50 μl ddH_2_O and stored at -80 °C. Samples were diluted 10x and enrichment over IN of the desired sequences was estimated with qPCR, using the iTaq Universal SYBR Green Supermix (Bio-Rad) with the indicated primers (**Table S3**). Data were plotted with GraphPad Prism 9.

### Chromatin Endogenous Cleavage assay (ChEC)

Samples and buffers were kept on ice at all times. Frozen fixed cell pellets were washed with 500 µl Buffer A (15 mM Tris-HCl pH = 7.4, 0.2 mM Spermine, 0.5 mM Spermidine, 80 mM KCl, pH = 8, 3 mM EDTA, 1x protease and phosphatase inhibitors) and then they were centrifuged at 20000 xg for 1:30 min at 4 °C. The supernatant was discarded and cell pellets were resuspended with 500 µl Buffer A. ∼500 μl precooled glass beads (1 mm) were added to cover the whole suspension. Cells were disrupted on a VXR basic IKA Vibrax orbital shaker at 2200 rpm at 4 °C for 15 min, three times with 10 min breaks on ice in between. The bottom and top of the 1.5 µL tube was pierced using a hot needle and placed into a 15 mL falcon tube. After centrifugation (130 xg, 2 min at 4 °C), the beads remained in the 1.5 μL tube and the lysate could be collected in the 15 mL tubes. The crude cell lysates were transferred into new 1.5 µl microtubes and centrifuged at 20000 g for 2 min at 4°C.

The supernatants were discarded and the pellets were resuspended in 500 µl Buffer A. After another centrifugation step (20000 × g for 2min at 4°C), supernatants were discarded and cell solutions were resuspended in 350 μl Buffer Ag (Buffer A without EDTA, with the addition of 0.1 mM EGTA) for a total volume of 400 - 450 μl. Samples were incubated for 3 min at 30° at 750 rpm. Then, samples were vortexed vigorously and 80 µl of suspensions were transferred to fresh DNA-low binding tubes containing 100 μl of IRN buffer, to be used as Controls (0 min ChEC). 7 μl of 0.1 M CaCl_2_ (2 μl per 100 μl of cell solution, final concentration 2 mM) was added in each tube and the samples were incubated at 30°C with constant shaking (750 rpm) for the activation of MNase digestion. After 10, 30, and 60 min of incubation, 80 µl of the samples were aliquoted into 1.5 μl DNA-low binding tubes containing 100 µl of IRN buffer and mixed to stop the MNase activity.

All samples were incubated with 4 µl of RNase A (10 mg/ml), mixed, and incubated at 37 °C for at least 1 hr. 10 µl of 10% SDS (final concentration 0.5%) and 4 µl of Proteinase K (10 mg/ml) were added, mixed, and samples were incubated for 1 hr at 56 °C. Crosslinking was reversed by incubation at 65 °C overnight. The next day, digested DNA was isolated by phenol-chloroform-isoamyl-alcohol (25:24:1) extraction. Purified DNA was washed with 70% EtOH and was resuspended in 25 μl ddH_2_O. 12 μl of each sample were transferred to fresh 1.5 μl tubes and were incubated with 2 μl Cutsmart buffer (NEB), 1.5 μl XmaI and PmlI restrictions enzymes (NEB) each (30 and 15 U, respectively) and 3 μl ddH_2_O (total volume 20 μl) at 37 °C overnight, to isolate the mAIRN reporter sequence (∼1.4 kb) for further detection. Next day, restricted samples were mixed with 4 μl Purple Loading Dye 6x (NEB) and were loaded on a 1% Agarose gel for the size-separation of the purified DNA and the digested and restricted DNA fragments. The agarose gel containing the separated DNA fragments was incubated with 0.2 N HCl for 20 min at RT. Then, it was rinsed with H_2_O, washed twice with Denaturation solution (0.5 M NaOH, 1.5 M NaCl) for 15 min at RT, rinsed once again with H_2_O, and washed twice with Transfer buffer (1 M NH_4_AcO).

### Southern Blot

For the Southern blot transfer assembly, a platform was placed in a tray containing transfer buffer (1 M NH_4_AcO). A wet Whatman paper, free of air bubbles, was placed on the platform. This paper served as a wick to draw the transfer buffer upwards through the gel. The UV-crosslinked gel was then carefully placed face-down on the wet Whatman paper. A pre-wet nylon membrane was positioned on top of the gel, ensuring no air bubbles were trapped. Three additional Whatman paper sheets were placed on the membrane, followed by a stack of paper towels and a light weight (approximately 0.5 kg). The entire assembly was left undisturbed overnight to allow complete transfer of the DNA fragments. Transfer times varied depending on the fragment size, with fragments up to 15 kb requiring approximately 18 hr. Following transfer, the membrane was UV crosslinked using the Stratalinker autocrosslinking function to covalently bind the DNA fragments, enhancing hybridization signals during subsequent detection steps. The membrane could then be dried and stored at room temperature for further analysis. The DNA probe (294 bp) detecting the mAIRN locus was generated by PCR using primers mAIRN-3-SB-Frw and mAIRN-3-SB-Rev from yeast genomic DNA from yeast strain mAIRN CoD as a template (**Tables S1&S3**). The resulting double-stranded PCR fragment was then used to generate the DNA probe by body labeling using the RadPrime DNA labeling system (Invitrogen) with incorporation of [α−32P] dATP (Hartmann Analytik) according to the instructions of the manufacturer. Membranes were prehybridized for 1 hr at hybridization temperature (65 °C) with 10-15 ml of hybridization buffer (2x SSC, 0.5 M sodium phosphate buffer pH 7.2, 7 % SDS). Following prehybridization, the buffer was discarded and replaced with 15 ml of fresh prewarmed hybridization buffer. The probe, mixed with salmon sperm DNA (final concentration 100 μg/ml) and boiled for 5 min, was then added to the tube. Hybridization occurred overnight at 65 °C with gentle rotation in a hybridization oven. Blots were washed once with 30 mL of 3x SSC and 0.1 % SDS after hybridization. Stringency washes were performed at hybridization temperature with rotation, using three buffers in sequential order: 0.3x SSC with 0.1 % SDS, 0.1x SSC with 0.1 % SDS, and lastly 0.1x SSC with 1.5 % SDS. Each wash step was repeated twice for 15 min. Finally, blots were dried, stored at room temperature, exposed to phosphorimaging screens and read out using a Typhoon scanner. Intensity values were normalized to the uncleaved mAIRN sequence (1.4 kb) intensity upon restriction of the ChEC-treated samples with XmaI and PmlI.

### RNA extraction, cDNA synthesis and quantitative RT-PCR

Yeast cells were grown to OD_600_ = 1 and 2 ml were harvested and centrifuged for 5 min. For the induction of PP7-mAIRN, cells were incubated with 2% GAL for 2 hr. Total RNA was isolated with the RNeasy Kit (QIAGEN) according to the manufacturer’s protocol, using zymolyase digestion (100U per sample). Purified RNA samples were incubated with DNase for the removal of DNA, according to the manufacturer’s protocol (QIAGEN). cDNA was synthesized with the SuperscriptIII First Strand Synthesis System (Invitrogen) according to the protocol, using poly-dT primers. Quantification of PP7-mAIRN transcription levels in different mutant strains and in different timepoints was conducted with quantitative RT-PCR using the Power SYBR Green PCR MasterMix (Thermo) according to the manufacturer’s protocol. Reactions were conducted with 0.4 ng of cDNA template, in triplicates for each combination of strain/set of primers. TDH3 or ACT1 genes were used as internal control using previously described primers [35]. The sequences of primers used for qRT-PCR are shown in **Table S3**. Data were plotted with GraphPad Prism 9.

### Quantification and statistical analysis

Time-lapse measurements were collected with ZEN3.0 and analyzed using a custom-made computational pipeline developed specifically for the analysis of replication rates and transcription dynamics during replisome progression by our group (Transcriptomatic5). Our MATLAB-based pipeline identifies, tracks and quantifies the LacI-Halo-SiR, tetR-tdTomato and PCP-Envy dots in each cell. Proper identification of the PCP-Envy dot that corresponds to the transcription site is validated by ensuring colocalization with the lacI-Halo-SiR and tetR-tdTomato dots, and the PCP-Envy back-ground signal is subtracted to quantify transcription site intensity. Quantification results for multiple cells in each strain are averaged, while normalizing the time axis for each individual cell according to its replication rate, such that the average transcriptional intensity is calculated relative to the location of the replisome along the chromosome. Statistical analysis of replication time data and single-cell transcription intensities was performed using Monte Carlo resampling with 1 000 000 iterations. Swarm plots and violin plots were plotted in Python.

## RESULTS

### Replication fork progression is not hindered by mAIRN R-loops in wildtype cells

Investigating the impact of R-loop formation on both replisome progression and transcription constitutes a challenge as it requires the simultaneous tracking of both processes at a specific genomic locus. Previously, we have developed such a system for monitoring transcription-replication coordination by combining locus-specific real-time detection of replisome progression and transcription bursting in single *Saccharomyces cerevisiae* cells by live-cell imaging [30]. This approach is based on tracking replisome progression through a single replicon in real-time [36]. The genomic locus of interest is labelled with tandem repeats of bacterial operator sequences, *lacO* and *tetO*, integrated in a defined and increasing distance from the early-efficient replication origin *ARS413*. These arrays are bound by their cognate repressors conjugated with fluorescent proteins; lacI fused to HaloTag, labelled with SiR-Halo dye (LacI-Halo-SiR) and tetR labelled with tdTomato (tetR-tdTomato) (**Fig. 1A**). This labelling enables the visualization of the chromosomal locus as two colocalizing fluorescent foci, measurement of *lacO/tetO* array duplication, and thus, replisome progression (**Fig. 1B**) [30,36].

**Figure 1.**
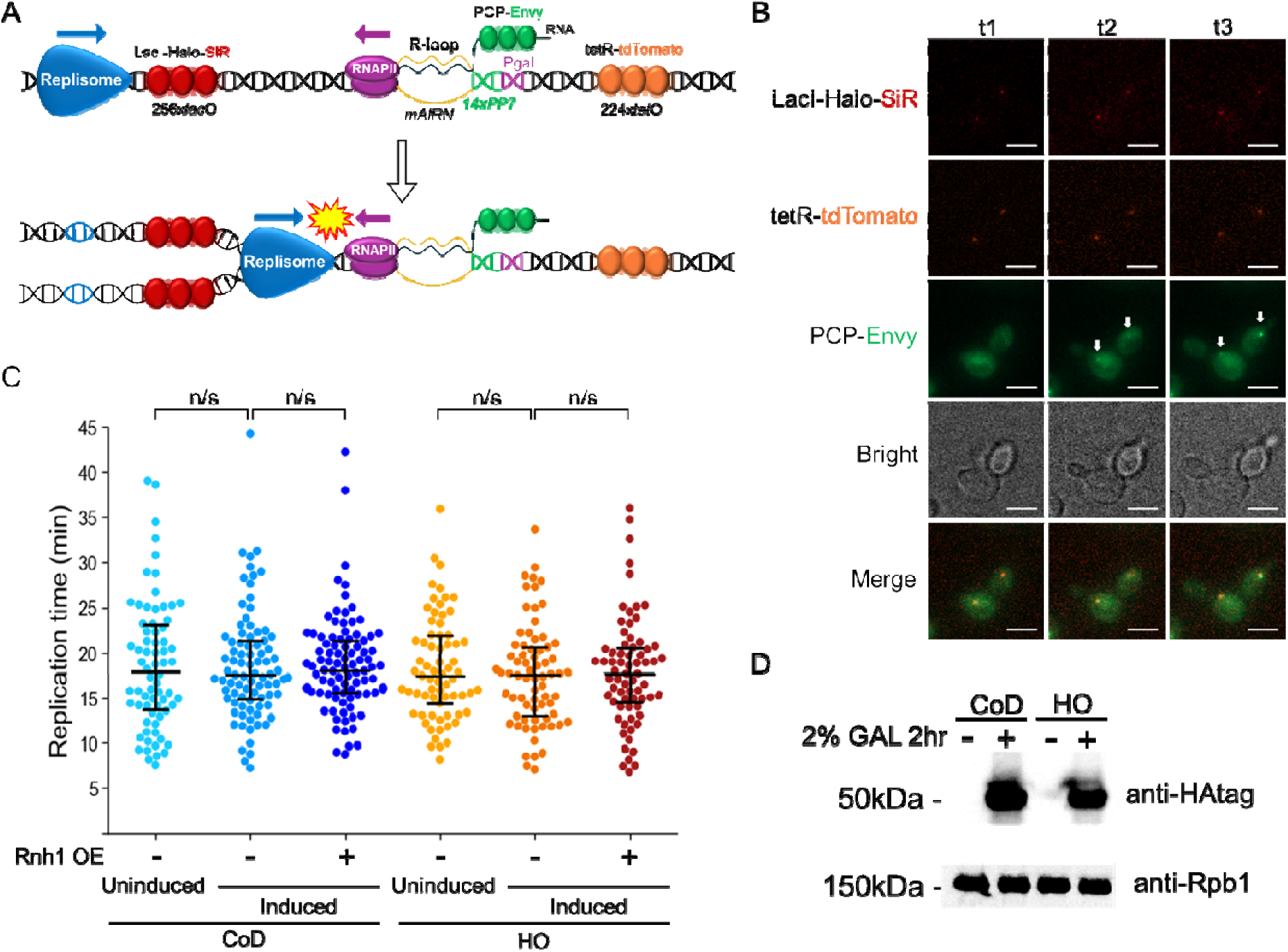
Transcription of mAIRN R-loop gene does not slowdown replication through the gene. (**A**) Schematic depiction of the live-cell imaging system for measuring replication times through the mAIRN sequence[37] (see text and [30] for details). (**B**) Representative cells with LacI-Halo-SiR and tetR-tdTomato-labeled *lacO* and *tetO* arrays, respectively, with PCP-Envy labelled mAIRN transcription bursts located in a CoD orientation. PCP-Envy foci, marking transcription bursts, colocalize with the two arrays (white arrows). Different cell cycle stages (t1 = G1-early S, t2 = Mid-late S, t3 = Late S-G2) are shown. Scale bar = 5 μm (**C**) Replication is not affected by R-loop presence in WT cells. Replication times of mAIRN-expressing cells in CoD and HO, in the absence or presence of induction, with (+) or without (-) Rnh1 OE. Median replication times and cell number for each yeast strain and condition are listed in **Table S6**. n/s = not significant (**D**) Representative Western Blot for the detection of Pgal-RNH1-3HA OE, in the presence or absence of galactose. Anti-Rpb1 is used as loading control.

To investigate how R-loop formation interferes with replisome progression in the labelled locus, we integrated a strong inducible R-loop-forming sequence in the path of the replisome (**Fig. 1A**). Previously, we devised a similar approach to induce a TRC, by integrating the *GAL10* gene under the control of its minimal native *GAL10* promoter (Pgal) [30] between the two bacterial operator arrays in both CoD and HO orientations. Here, we replaced the *GAL10* open reading frame (GAL10orf) with the R-loop-forming sequence of mAIRN [37]. This mAIRN sequence has been previously shown to generate stable and persistent R-loops both *in vitro* and *in vivo*, and it has been utilized for studying the effects of R-loops on replication fork speed, transcription competence, and DNA damage/signalling [20,38–43],[44] (**Fig. 1**).

Here, using this system, we measured changes in mAIRN transcription during replisome passage through the transcribing mAIRN gene. To monitor transcription during cell cycle progression, we labelled the 5’-end of mAIRN with 14 repeats of the PP7 bacteriophage sequence in cells expressing the bacteriophage PP7 coat protein fused to an optimized GFP (PCP-Envy) [30,45] (**Fig. 1A&B**).

First, we examined replication fork progression in the presence or absence of mAIRN transcription. Surprisingly, we did not detect significant replication slowdown, neither when mAIRN was transcribed in CoD, nor in HO orientation (**Fig. 1C** and **Table S6)**. In accordance, Rnh1 overexpression (OE) did not affect replication rates (**Fig. 1C&D** and **Table S6**). These results indicate that in WT cells, replication is robust even during passage through a strong R-loop forming sequence.

### R-loops repress mAIRN transcription in a replication dependent manner

To monitor mAIRN transcription dynamics before, during and following replisome progression through the mAIRN gene, we normalized and aligned replication and transcription measurements of all individual cells as previously described [30] (**Fig. 2**). Strikingly, mAIRN transcription in G1 and early S phase cells, in both CoD and HO orientations, was surprisingly low, close to background levels (**Fig. 2A&B** and **Fig. S1A&B**). Notably, upon replication fork passage through the mAIRN sequence, we detected a dramatic increase of ∼2-fold (100%) of transcription intensity compared to the transcription level before mAIRN replication (**Fig. 2A&B & S1A&B).** This transcriptional increase occurred approximately 9 kb after the replication fork passed the mAIRN gene (**Table S7**), indicating a fast activation of gene expression following mAIRN replication, consistent with previous reports [30,46]. The same analysis for the yeast strains expressing the non R-loop forming PP7-labelled *GAL10* showed only ∼1.2-fold (20%) increase after replisome passage ([30] and **Fig. S1C**), suggesting that this strong transcriptional upregulation is specific to the R-loop forming mAIRN sequence.

**Figure 2.**
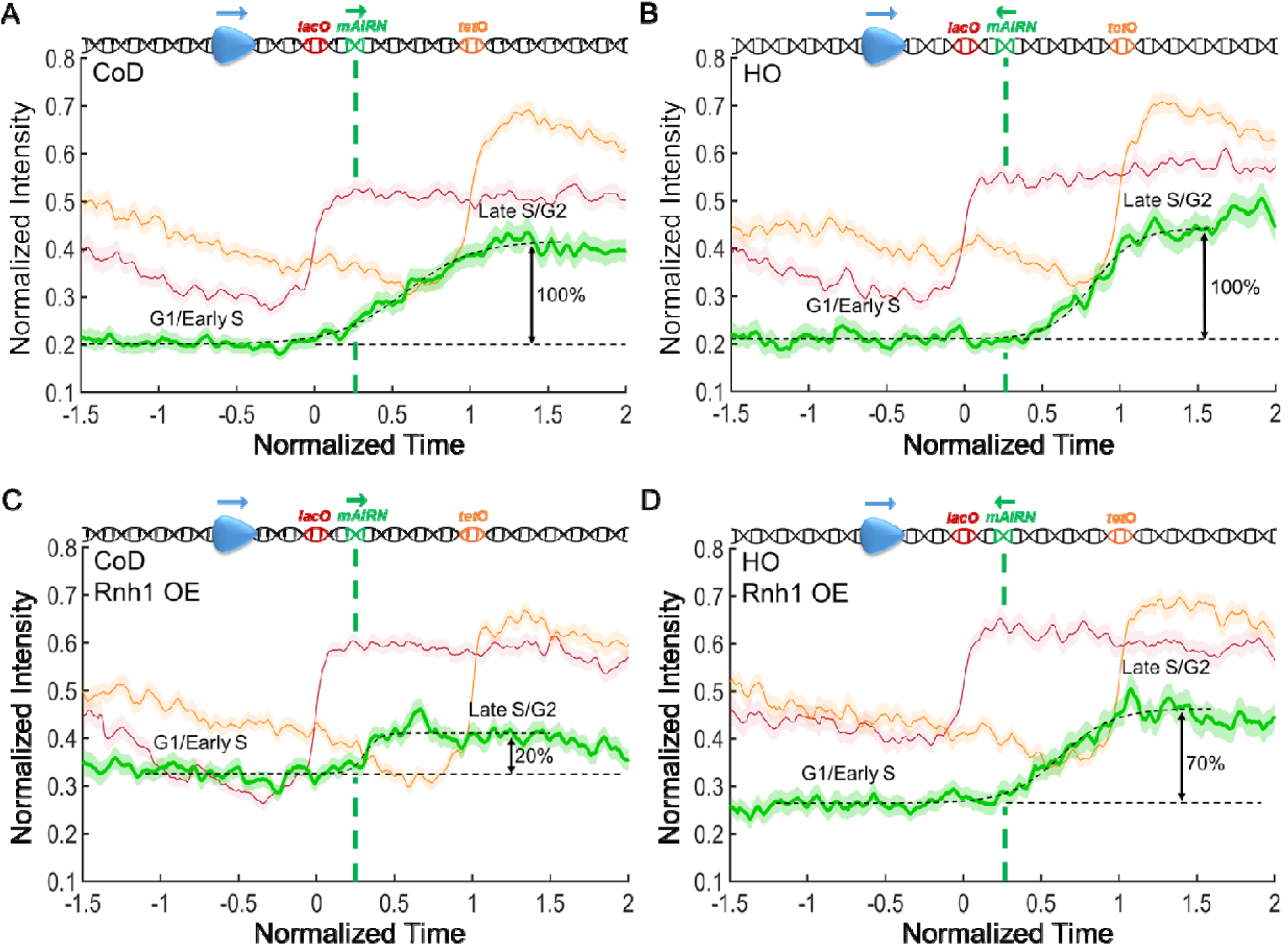
Replication-dependent R-loop repression of mAIRN transcription. (**A-D**) Average normalized fluorescent intensity of the transcription site (PCP-Envy focus, green) and the labelled *lacO* and *tetO* arrays (LacI-Halo-SiR and tetR-tdTomaro foci, red and orange respectively) were plotted over time for cells transcribing mAIRN (Induced) in (**A** and **C**) CoD or (**B** and **D**) HO orientation relative to the replisome originating from *ARS413*, in (**A** and **B**) WT or (**C** and **D**) Rnh1 OE conditions. The time axis is normalized according to the duplication times of the arrays. Shaded areas represent the standard error of the mean (SEM). A green dashed vertical line represents mAIRN duplication time. Dashed black lines represent a fit of the transcription intensity data to a sigmoidal function (see **Table S7** for parameters). Top: Scheme showing the relative positions of *ARS413*, *lacO* and *tetO* arrays, and mAIRN relative to replisome movement on this locus.

It has been shown that following gene replication, cells maintain stable transcript levels during G2, through a process called transcription homeostasis [47]. As such homeostasis is not observed following mAIRN duplication, we hypothesized that its transcription is suppressed before replication, possibly as a result of R-loop formation.

To test this hypothesis, we overexpressed Rnh1 in our strains to deplete mAIRN R-loops. Interestingly, we found that Rnh1 overexpression (OE) significantly decreased the difference in transcription between early and late S phase, most likely due to increased mAIRN transcription in G1/early S phase cells (**Fig. 2C&D** and **S1D-G**). These results suggest that R-loops repress mAIRN transcription prior to gene replication. This phenotype was more prominent in the case of CoD mAIRN transcribing strains (**Fig. 2C** and **S1D&F**). Importantly, transcription was not altered when we overexpressed Rnh1 in the yeast strains expressing PP7-*GAL10* ([30], **Fig. S1H**), further supporting the R-loop dependency of these effects. Notably, we were unsuccessful in the direct detection of the differences in R-loop levels at the mAIRN gene under these conditions by bulk DRIP-qPCR assays, likely due to cell-to-cell variations in S phase entry and cell cycle progression.

Taken together, our results indicate that R-loop formation represses mAIRN transcription prior to gene replication. However, following gene replication mAIRN transcription is upregulated, highlighting replisome progression through the mAIRN as a key regulatory process in R-loop mediated gene repression.

### R-loops inhibit RNAPII activity in a replication dependent manner

To investigate whether elevated mAIRN transcription upon R-loop removal is mediated by increased RNAPII occupancy, we examined RNAPII binding to mAIRN using RNAPII Chromatin Immunoprecipitation followed by quantitative PCR (ChIP-qPCR) analysis. In agreement with the live-cell imaging experiments, Rnh1 OE resulted in increased levels of RNAPII binding on mAIRN, both in 5’ and 3’ ends of CoD and HO yeast strains (**Fig. 3A**). Importantly, such RNAPII enrichment upon Rnh1 OE was not detected in the highly expressed ACT1 housekeeping gene (**Fig. S2A**).

**Figure 3.**
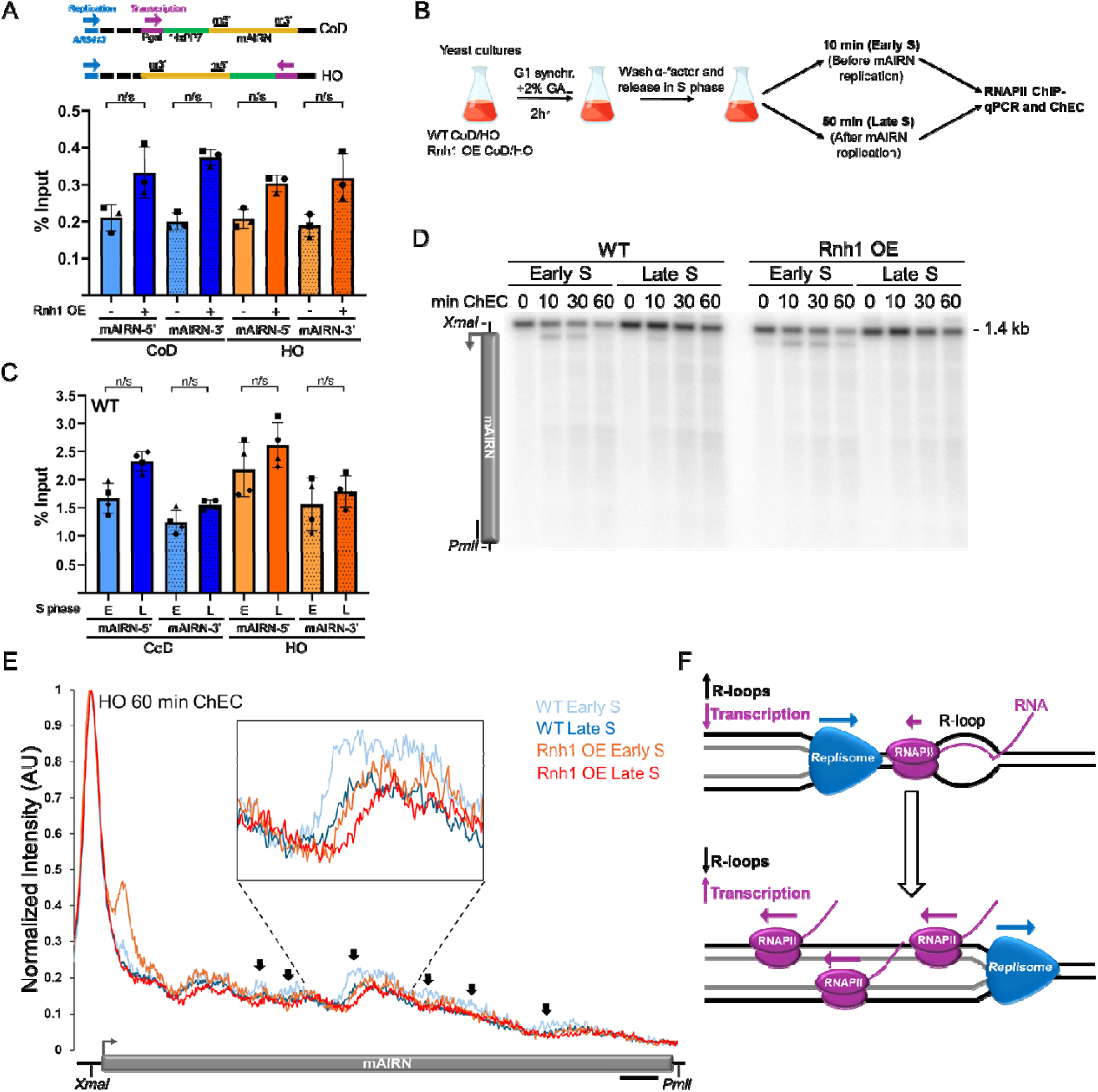
R-loops limit RNAPII binding and processivity of mAIRN sequence. (A) RNAPII ChIP-qPCR analysis of mAIRN CoD and HO-expressing strains shows elevated RNAPII occupancy upon Rnh1 OE. % Input values were normalized with TDH3. Averages of three biological replicates are shown. Error bars represent ±SD. (B) Experimental strategy to investigate how replisome progression affects RNAPII occupancy and activity on mAIRN. (**C**) RNAPII ChIP-qPCR analysis of WT CoD and HO strains using the two primer sets used in (**A**) shows elevated RNAPII occupancy in Late S phase timepoint (post-mAIRN replication). % Input values were normalized with ACT1 values. The mean of four biological replicates is shown. Error bars represent ±SD. (**D**-**E**) ChEC analysis of RNAPII occupancy throughout HO mAIRN cells, sampled as in (**B**) (early or late S phase), with or without Rnh1 OE. (**D**) Representative Southern Blot membrane of restricted ChEC-treated samples (0, 10, 30, 60 min). Left: Scheme of mAIRN sequence with marked restriction sites and probe location aligned with the Southern Blot analysis. (**E**) Relative RNAPII occupancy through mAIRN in the HO orientation yeast strain, both in WT (light and dark blue) and Rnh1 OE (orange and red) cells, as measured with ChEC-Southern blot upon 60 min MNase digestion. Black arrows pinpoint mAIRN locations where differences in RNAPII occupancy were detected. Magnification of one of these regions is presented. One of two biological replicates is shown. Bottom: Scheme of mAIRN sequence aligned with the relative RNAPII occupancy quantification plot. (**F**) Suggested mechanism for replisome-dependent mAIRN transcription derepression. n/s = not significant

Next, we examined whether RNAPII binding to mAIRN changes during cell cycle progression. Specifically, we sampled cells 10 min and 50 min post G1 release to test RNAPII occupancy before and after mAIRN replication, respectively (**Fig. 3B**). As expected, we detected higher RNAPII abundance at the 5’ compared to 3’-end of the mAIRN sequence in both CoD and HO strains (**Fig. 3C**), consistent with the typical intragenic distribution of RNAPII on actively transcribed genes [48]. Furthermore, we detected a 1.5 – 2-fold higher RNAPII occupancy on mAIRN gene body in late S compared to early S phase cells, in both CoD and HO mAIRN-expressing yeast strains (**Fig. 3C**). Surprisingly, however, Rnh1 OE did not affect this pattern (**Fig. S2B**), despite a major transcriptional upregulation observed in G1/early S phase cells by live cell microscopy (**Fig. 2C&D**). This data suggests that the observed downregulation of transcription prior to replication was not caused exclusively by reduced RNAPII binding on the mAIRN gene but could rather be explained by R-loop dependent defects in RNAPII elongation.

Subsequently, we tested whether particularly strong RNAPII pausing sites can be detected within the mAIRN sequence upon R-loop formation. To this end, we fused the second-largest RNAPII subunit, RPB2, with Micrococcal nuclease (MNase) to its C-terminal end (**Fig. S2C**) and performed Chromatin Endonuclease Cleavage (ChEC) assay followed by Southern blot analysis. The cell cycle dependent ChEC assay (**Fig. 3B**) revealed that, in both CoD and HO orientations, RNAPII accumulates on specific regions at about 500-700 bp in the centre of the mAIRN sequence (**Fig 3D&E**, black arrows, and **S2D&E**). Interestingly, this accumulation of RNAPII pausing sites appears to be more pronounced before replisome progression compared to after replisome passage and is strongly reduced by Rnh1 OE (**Fig. 3E** and **S2E**). Overall, these results show that R-loops inhibit RNAPII binding and mobility during mAIRN transcription and replication fork progression can overcome these transcriptional defects (**Fig. 3F**).

### RnaseH enzymes are essential for replication and transcription at the mAIRN gene

To examine the roles of RNHs in coordinating TRCs at R-loops, we deleted both catalytic subunits of the two main *S. cerevisiae* RNHs, *rnh1*Δ and *rnh201*Δ (RNH1 and RNH2, respectively). It has been thoroughly reported that the absence of RNHs leads to extensive, genome-wide R-loop formation, R-loop-dependent replisome stalling, and elevated levels of genome instability [37–40]. Consistent with these studies, replication time of induced RNH mutant cells was significantly longer compared to the uninduced condition, in both CoD and HO orientations (**Fig. 4A** and **Table S6**). Importantly, Rnh1 OE in these strains restored normal replication fork progression following mAIRN induction (**Fig. 4A** and **Table S6**). These results highlight that the two RNHs are indispensable for the removal of excessive R-loops and their action is a prerequisite for normal replisome progression through mAIRN R-loops, in agreement with previous research [49].

**Figure 4.**
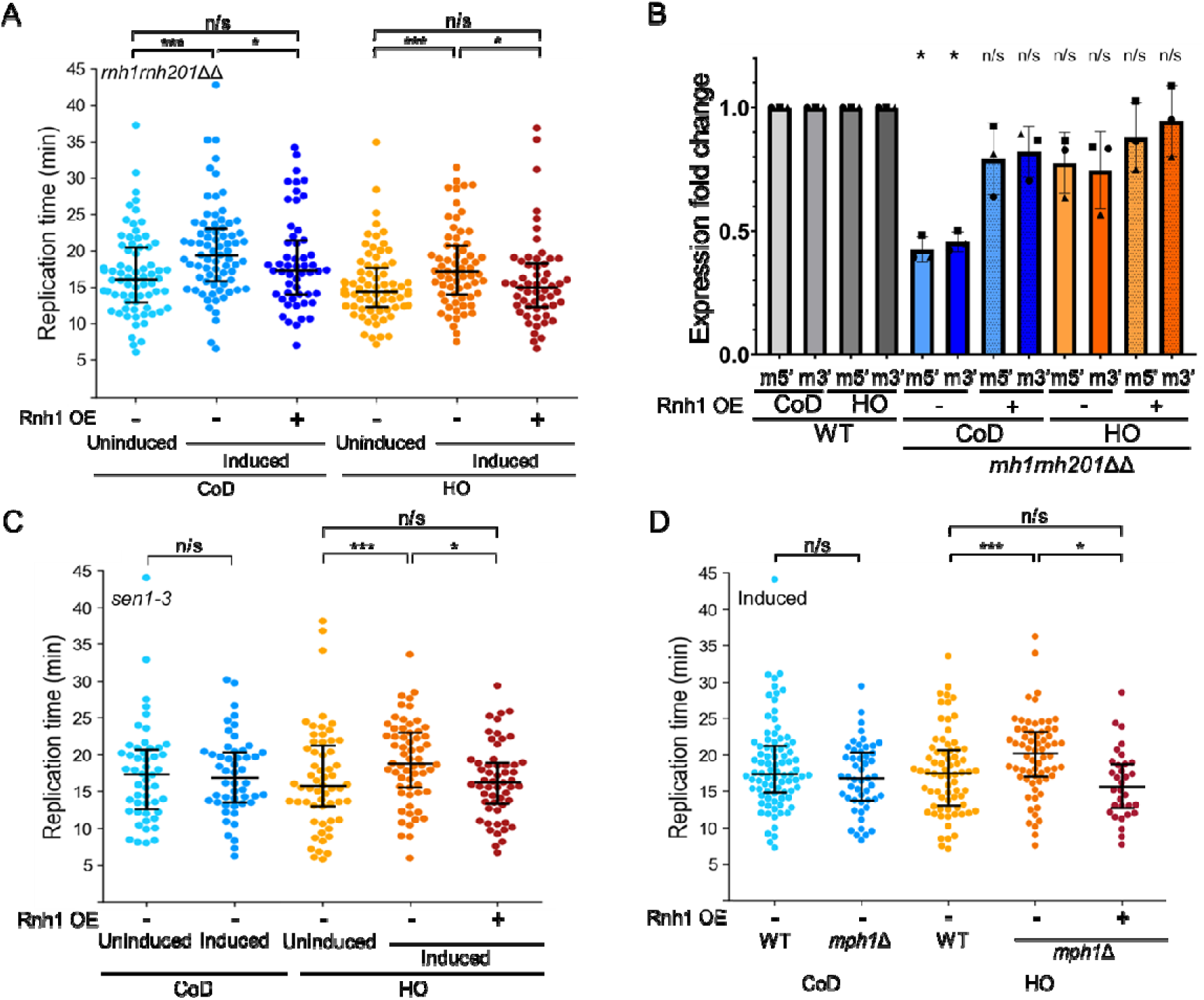
R-loops in mutant yeast strains slowdown replisome progression and can decrease mAIRN transcription. (**A**) Replication is slowed down in *rnh1rnh201*ΔΔ strain upon mAIRN induction in an R-loop dependent manner. Replication times in CoD and HO *rnh1rnh201*ΔΔ cells, in the absence or presence of induction, with (+) or without (-) Rnh1 OE. (**B**) RT-qPCR analysis of mAIRN expression levels in induced WT or *rnh1rnh201*ΔΔ cells, with or without Rnh1 OE, using the primer sets shown in Fig. 3A. The average of three biological replicates is shown. Error bars represent ±SD. (**C-D**) Activation of HO mAIRN transcription leads to replication slowdown in *sen1-3* and *mph1*Δ cells in an R-loop-dependent manner. Replication times as shown as in in (**A**), but for (**C**) *sen1-3* or (**D**) *mph1*Δ cells. WT replication times are adapted from Fig. 1C and are shown here for comparison. Median replication times and cell numbers for each yeast strain and condition are listed in **Table S6**. * = p < 0.05, *** = p < 0.005, n/s = not significant.

Next, we monitored mAIRN transcription during replication fork progression in the *rnh1rnh201*ΔΔ strains. Transcription in these strains show similar dynamics as WT, where transcription levels are very low before mAIRN replication but is strongly elevated upon replication fork passage through mAIRN (**Fig. S3A&B**). Rnh1 OE did not alter these profiles, although interestingly, the increase of PCP-Envy foci intensity after replisome passage from mAIRN was more gradual than without Rnh1 OE, indicating that R-loops may affect the kinetics of transcription recovery following gene replication (**Fig. S3A-D** and **Table S7**).

Next, we tested whether excessive R-loop levels in the case of *rnh1rnh201*ΔΔ could affect the absolute mAIRN transcription levels using RT-qPCR. In accordance with live-cell experiments, mAIRN RNA levels in the double mutant diminished compared to WT cells (**Fig. 4B**). Specifically, transcription levels of CoD mAIRN were reduced 50-60% compared to WT. Strikingly, Rnh1 OE restored RNA levels almost to WT (**Fig. 4B**), further demonstrating the repressive effect of R-loops on mAIRN transcription.

### Sen1 and Mph1 are essential for normal mAIRN replication without affecting transcription

RNA:DNA helicases, such as Sen1 and Mph1, have been previously shown to safeguard the genome from R-loop-dependent replisome stalling and genomic instability [15,50–55]. To study the importance of Sen1 and Mph1 for coordinating TRCs at R-loops, we performed live-cell microscopy experiments in *sen1* and *mph1* mutants. In the case of Sen1, we focused on its interaction with the replication fork, and thus we generated the Sen1 replisome binding-deficient mutant (*sen1-3*) [32]. Microscopy analysis revealed that both *sen1-3* and *mph1*Δ mutations slowdown replisome progression through the HO transcribing mAIRN gene but have no significant effect when mAIRN is in CoD orientation (**Fig. 4C&D** and **Table S6**). Rnh1 OE in these strains restored normal replication through the transcribing mAIRN gene, indicating the harmful effect of HO R-loops on replisome progression in the absence of these helicases (**Fig. 4C** and **Table S6**).

The mAIRN transcriptional patterns of *sen1*-*3* or *mph1*Δ mutant cells were not significantly altered compared to WT cells (**Fig. S4**). Nevertheless, upon Rnh1 OE, we noticed a slight elevation of transcription levels in the HO yeast mutant strains, specifically before replisome passage (**Fig. S4,** G1/Early S), demonstrating that R-loop depletion can upregulate mAIRN transcription even in RNA:DNA helicase-mutant strains.

### The balance between RNAPII binding and R-loop formation prevents harmful TRCs

We were intrigued by the mechanism of harmful TRCs associated with excessive or unregulated R-loops, leading to replisome stalling. Several studies have previously examined whether the RNA:DNA hybrids themselves, the bound RNAPIIs, or both are the source of R-loop dependent TRCs [56]. In order to investigate the interplay between RNAPII binding and R-loop formation, we generated and examined the *rpb1-1* mutant on the background of the mAIRN strain in the HO orientation. This mutant was shown to have a higher affinity for the DNA [57] and in our previous work, we found that it leads to replication slowdown upon induced TRCs at the non-R-loop *GAL10* gene [30]. Initially, we performed RNAPII ChIP-qPCR experiments to examine the RNAPII retention levels in the *rpb1-1* mutant strains in the absence or presence of Rnh1 OE. We found that Rnh1 OE leads to a dramatic and specific increase in RNAPII retention on the mAIRN sequence (**Fig. 5A** and **Fig. S5A**), in agreement with the inhibitory effect of R-loops on mAIRN transcription.

**Figure 5.**
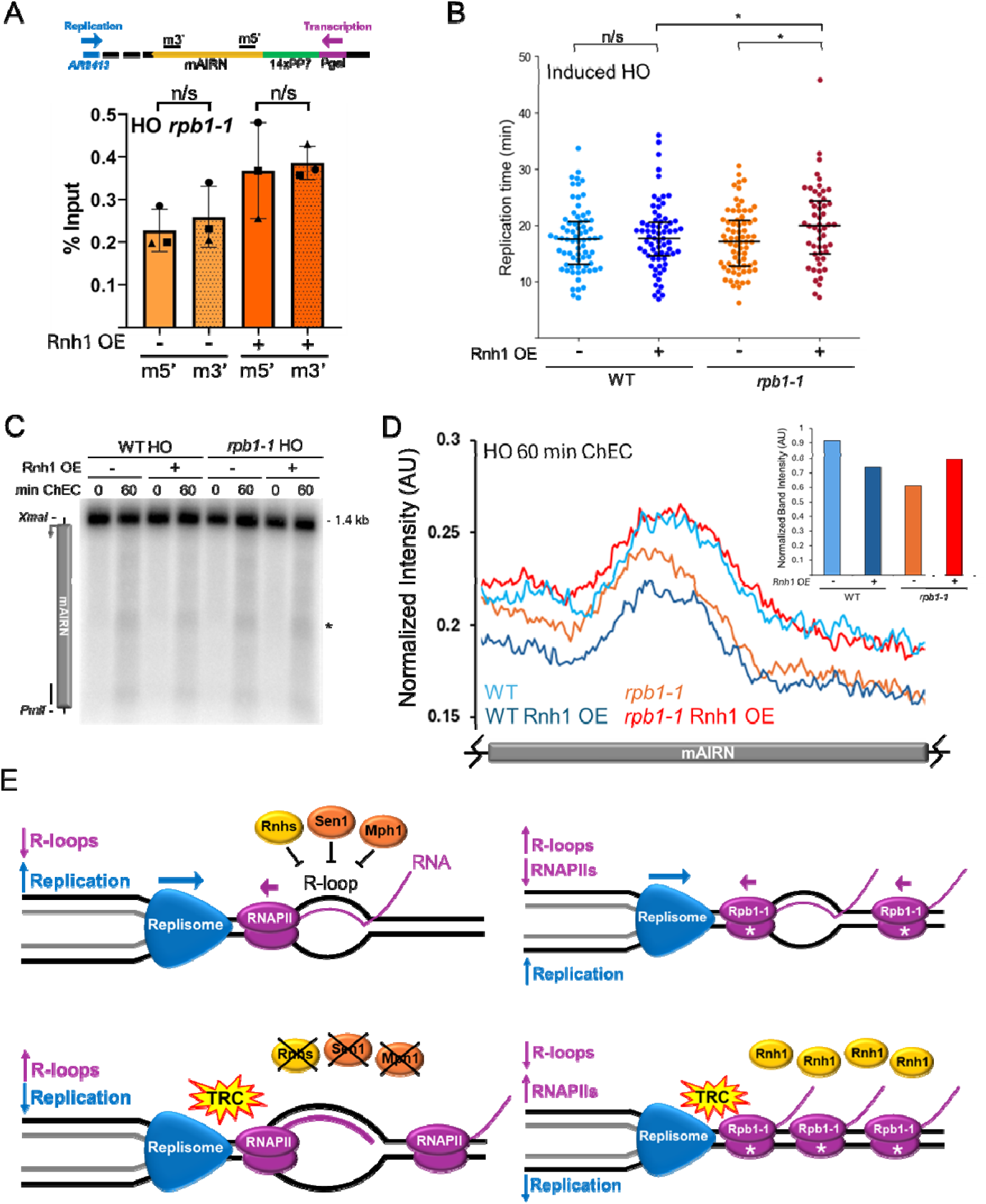
R-loops regulate RNAPII binding and mobility and prevent replication slowdown in *rpb1-1* mutant cells. **(A)** Elevated RNAPII occupancy upon Rnh1 OE detected by ChIP-qPCR analysis of *rpb1-1* mAIRN HO-expressing strains using the primer sets shown in Fig. 3A. **(B)** Slowdown of replication in *rpb1-1* cells upon mAIRN induction and Rnh1 OE (**Table S6**). WT replication times are adapted from Fig. 1C and are shown here for comparison. * = p < 0.05, n/s = not significant. (**C-D**) ChEC analysis of RNAPII occupancy throughout HO mAIRN, in WT or *rpb1-1* cells, with or without Rnh1 OE shows the increased binding of RNAPII in *rpb1-1* cells upon Rnh1 OE. (**C**) Southern Blot membrane of ChEC-treated samples (0 and 60 min) for the different strains (as described in Fig. 3C). An asterisk depicts the region/band quantified in (**D**). (**D**) Relative RNAPII occupancy and pausing on the mAIRN region within a specific region in mAIRN (see **Fig. S5E**), in WT or *rpb1-1* cells, with or without Rnh1 OE, as quantified by Southern Blot. The top right graph depicts the quantification of the band labeled with an asterisk in (**C**). One of two biological replicates are shown. Bottom: scheme of mAIRN sequence aligned with the relative RNAPII quantification plot. (**E**) Two independent mechanisms for TRC leading to fork slowdown. Left: R-loop-formation leads to transcriptional repression allowing unperturbed replication. Mutations in RNHs or RNA:DNA helicases lead to harmful TRCs leading to R-loop dependent replication slowdown. Right: R-loops regulate RNAPII binding to mAIRN for *rpb1-1* allowing unperturbed replication. However, R-loops removal leads to excess RNAPIIs binding and replication slowdown.

Next, we examined replication fork progression in the *rpb1-1* mutant in the presence or absence of Rnh1 OE upon mAIRN induction. Interestingly, in the absence of Rnh1 OE, we found that replication is not affected in *rpb1-1* cells, indicating that R-loop formation at the mAIRN gene may prevent excessive RNAPII binding in these mutant cells (**Fig. 5B** and **Table S6**). However, in the presence of Rnh1 OE, we observed replication slowdown in agreement with enhanced RNAPII retention on mAIRN (**Fig. 5A**) and previous measurements with induced *GAL10* gene [30]. Such high RNAPII retention upon Rnh1 OE also led to elevation of transcription levels during early S phase (**Fig. S5B-D**, G1/Early S transition).

Finally, we performed ChEC experiments to reveal changes of RNAPII occupancy in *rpb1-1* mutant cells and detect pausing sites of RNAPII along the mAIRN gene. While *rpb1-1* had minor effects on RNAPII distribution through the mAIRN gene, Rnh1 OE resulted in a significant increase of RNAPII coverage, both in specific hotspots and spanning the whole mAIRN sequence (**Fig. 5C&D** and **S5E-H**).

Overall, our results with *rpb1-1* mutant strains indicate an unexpected role for R-loops in reducing excessive RNAPII occupancy on the gene, preventing replisome stalling and harmful TRCs. Thus, our study highlights that “trains” of multiple RNAPIIs occupying short stretches of a highly transcribed gene are the more potent obstacle to replisome progression than when facing lower levels of RNAPII molecules at R-loops.

## DISCUSSION

R-loop formation has been linked with adverse consequences on DNA replication and transcription, giving rise to DNA damage and mutagenesis. In the current study, we directly examined the effects of R-loops on replication fork progression and explored the interplay between R-loops and transcription during replisome progression, in WT and mutant yeast strains.

### R-loop-regulation prevents harmful TRCs

Our observations in the current work are based on employing the well characterized mAIRN R-loop-forming sequence, for a mechanistic study in yeast. By measuring single-cell replication rates, we found that replisome progression is not affected by mAIRN R-loops, in either directionality (HO or CoD) (**Fig. 1C**). These results highlight the robustness of eukaryotic replisomes and the importance of multiple R-loop restricting mechanisms for maintaining normal replication even through R-loops. Indeed, only when the catalytic subunits of RNHs, *rnh1* and *rnh201*, were deleted, we observed significant replisome stalling, indicating the harmful effects of persistent mAIRN R-loops (**Fig. 4A**). Moreover, replication in the helicase-mutant yeast strains (*sen1-3* and *mph1*Δ) was also slowed down in the presence of HO R-loops (**Fig. 4C&D**). These findings are in agreement with several reports [20,44,57–62], underlying the importance of R-loop regulation in the HO orientation for maintaining genome stability.

### R-loop-mediated transcriptional repression is replication dependent

Our findings that R-loops repress mAIRN transcription are in accordance with previous reports showing that RNA molecules hybridized with the complementary DNA strand are an obstacle to transcription elongation, both *in vivo* and *in vitro* [22,23,60,63–66]. More recently, Mérida-Cerro et al. also showed transcription propagation impairment by expressing another heterologous R-loop prone sequence (Sμ350) in yeast cells [25]. Surprisingly, our finding that transcriptional repression of mAIRN occurs only prior to gene replication and can be rescued by Rnh1 OE reveals the dynamic crosstalk between replisome progression and mAIRN transcription regulated by R-loop formation (**Fig. 2**).

The R-loop mediated transcription repression can be caused by RNAPII eviction and/or impaired processivity, specifically before mAIRN sequence replication (**Fig. 3C-F**), suggesting the existence of R-loop removal mechanisms during gene replication. Such replication-dependent R-loop removal could be explained by the recruitment of R-loop-repressive factors by the replisome to the R-loop site. Indeed, both RNHs and Sen1 have been shown to interact with replisome components, highlighting their possible role in limiting R-loop formation during active DNA replication [67,68], facilitating both replication and transcription during transcription-replication encounters. Whether Mph1 also directly interacts with the replisome remains to be studied.

Alternatively, accumulated topological stress downstream of the replication fork could also contribute to the inhibition of R-loop formation. Another possibility is that chromatin changes during DNA replication, such as histone modifications and nucleosome remodelling or deposition, can negatively influence R-loop formation during gene replication [44,61,69–71]. Finally, direct dissociation of the RNA molecules from the complementary DNA strand, induced by the replicative CMG helicase traveling on the front of the leading strand [72] or FACT complex [61], can also be important. Different mechanisms for controlling R-loop levels during replication of CoD and HO genes may exist counting for the differences in transcriptional repression and recovery during CoD or HO mAIRN replication (**Fig. 2C&D** and **Fig. S1D-G**) [20].

### R-loops regulate RNAPII mAIRN binding to prevent replisome stalling

Our findings that R-loop depletion by Rnh1 OE in *rpb1-1* mutant strain leads to excessive RNAPII binding and replisome delays (**Fig. 5A-D**) highlight an unexpected beneficial role of R-loops in preventing TRCs. Overall, our results reveal two possible independent causes for harmful TRCs leading to replication fork slowdown: (i) accumulation of harmful R-loops in the absence of RNHs or RNA:DNA helicases (**Fig. 5E**, left) and (ii) accumulation of RNAPII on the transcribed gene (**Fig. 5E**, right). In the latter case, the presence of R-loops prevents excessive RNAPII binding, minimizing the possibility of direct collisions between the replisome and RNAPII (**Fig. 5E**, right). In these cases, functional R-loop-removal machinery, including RNHs and helicases, can ensure that the R-loops themselves will not stall replisome progression. Thus, a regulated level of R-loops may be beneficial at highly expressed genes to reduce the relative RNAPII load. As R-loops have been thoroughly reported to be enriched in highly-expressed genes [73,74], our results that R-loops regulate RNAPII occupancy at these genes may showcase the beneficial role of R-loops to allow replisome progression and prevent harmful TRCs.

## DATA AVAILABILITY

The code used for the analysis of live-cell imaging experiments is available online (**Table S5**). Any additional information required to reanalyze the data reported in this paper is available from the lead contact upon request.

## SUPPLEMENTARY DATA

Supplementary data is available at NAR online

## AUTHOR CONTRIBUTIONS

Conceptualization: S.H., A.A., I.T.

Data Curation: I.T.

Formal Analysis: I.T.

Funding acquisition: S.H. and A.A.

Investigation: Everyone

Supervision: S.H. and A.A.

Validation: Everyone

Writing – original draft: I.T.

Writing – review and editing: Everyone

## ACKNOWLEDGMENTS

We thank all past and current Hamperl lab and Aharoni lab members. We thank Philippe Pasero, Jerome Poli, and Jonathan Heuze for discussions and their critical assistance in DRIP assays. We thank Tineke Lenstra and Wim Pomp for advice and feedback in singe-cell transcription analysis. We thank Aziz El Hage for discussions and advice on DRIP assay. Ioannis wants to thank his wife, his family, and his cat.

## References

1. Rondón AG, Aguilera A. What causes an RNA-DNA hybrid to compromise genome integrity? DNA Repair (Amst) 2019;81:102660.

2. Petermann E, Lan L, Zou L. Sources, resolution and physiological relevance of R-loops and RNA–DNA hybrids. Nat Rev Mol Cell Biol 2022;23:521–40.

3. Hegazy YA, Fernando CM, Tran EJ. The balancing act of R-loop biology: The good, the bad, and the ugly. J Biol Chem 2020;295:905–13.

4. Mackay RP, Xu Q, Weinberger PM. R-Loop Physiology and Pathology: A Brief Review. DNA Cell Biol 2020;39:1914–25.

5. Brickner JR, Garzon JL, Cimprich KA. Walking a tightrope: The complex balancing act of R-loops in genome stability. Mol Cell 2022;82:2267–97.

6. Crossley MP, Song C, Bocek MJ et al. R-loop-derived cytoplasmic RNA-DNA hybrids activate an immune response. Nature 2023;613:187–94.

7. Wahba L, Costantino L, Tan FJ et al. S1-DRIP-seq identifies high expression and polyA tracts as major contributors to R-loop formation. Genes Dev 2016;30:1327–38.

8. Chan YA, Aristizabal MJ, Lu PYT et al. Genome-Wide Profiling of Yeast DNA:RNA Hybrid Prone Sites with DRIP-Chip. PLoS Genet 2014;10, DOI: 10.1371/journal.pgen.1004288.

9. Stirling PC, Chan YA, Minaker SW et al. R-loop-mediated genome instability in mRNA cleavage and polyadenylation mutants. Genes Dev 2012;26:163–75.

10. Brambati A, Colosio A, Zardoni L et al. Replication and transcription on a collision course: Eukaryotic regulation mechanisms and implications for DNA stability. Front Genet 2015;6:1–8.

11. García-Muse T, Aguilera A. Transcription-replication conflicts: How they occur and how they are resolved. Nat Rev Mol Cell Biol 2016;17:553–63.

12. Hamperl S, Cimprich KA. Conflict Resolution in the Genome: How Transcription and Replication Make It Work. Cell 2016;167:1455–67.

13. Wei X, Samarabandu J, Devdhar RS et al. Segregation of transcription and replication sites into higher order domains. Science 1998;281:1502–6.

14. Azvolinsky A, Giresi PG, Lieb JD et al. Highly Transcribed RNA Polymerase II Genes Are Impediments to Replication Fork Progression in Saccharomyces cerevisiae. Mol Cell 2009;34:722–34.

15. Aiello U, Challal D, Wentzinger G et al. Sen1 is a key regulator of transcription-driven conflicts. Mol Cell 2022;82:2952–2966.e6.

16. Skourti-Stathaki K, Proudfoot NJ. A double-edged sword: R loops as threats to genome integrity and powerful regulators of gene expression. Genes Dev 2014;28:1384–96.

17. Sollier J, Cimprich KA. Breaking bad: R-loops and genome integrity. Trends Cell Biol 2015;25:514–22.

18. Lalonde M, Trauner M, Werner M et al. Consequences and resolution of transcription–replication conflicts. Life 2021;11, DOI: 10.3390/life11070637.

19. Zardoni L, Nardini E, Brambati A et al. Elongating RNA polymerase II and RNA:DNA hybrids hinder fork progression and gene expression at sites of head-on replication-transcription collisions. Nucleic Acids Res 2021;49:12769–84.

20. Hamperl S, Bocek MJ, Saldivar JC et al. Transcription-Replication Conflict Orientation Modulates R-Loop Levels and Activates Distinct DNA Damage Responses. Cell 2017;170:774–786.e19.

21. Meng Y, Zou L. Building an integrated view of R-loops, transcription, and chromatin. DNA Repair (Amst) 2025;149:103832.

22. Tous C, Aguilera A. Impairment of transcription elongation by R-loops in vitro. Biochem Biophys Res Commun 2007;360:428–32.

23. Huertas P, Aguilera A. Cotranscriptionally formed DNA:RNA hybrids mediate transcription elongation impairment and transcription-associated recombination. Mol Cell 2003;12:711–21.

24. Poli J, Gerhold CB, Tosi A et al. Mec1, INO80, and the PAF1 complex cooperate to limit transcription replication conflicts through RNAPII removal during replication stress. Genes Dev 2016;30:337–54.

25. Mérida-Cerro JA, Maraver-Cárdenas P, Rondón AG et al. Rat1 promotes premature transcription termination at R-loops. Nucleic Acids Res 2024;52:3623–35.

26. Ginno PA, Lim YW, Lott PL et al. GC skew at the 5’ and 3’ ends of human genes links R-loop formation to epigenetic regulation and transcription termination. Genome Res 2013;23:1590–600.

27. Cerritelli SM, Crouch RJ. Ribonuclease H: the enzymes in eukaryotes. FEBS J 2009;276:1494–505.

28. Yang S, Winstone L, Mondal S et al. Helicases in R-loop Formation and Resolution. J Biol Chem 2023;299:105307.

29. García-Muse T, Aguilera A. R Loops: From Physiological to Pathological Roles. Cell 2019;179:604–18.

30. Tsirkas I, Dovrat D, Thangaraj M et al. Transcription-replication coordination revealed in single live cells. Nucleic Acids Res 2022, DOI: 10.1093/nar/gkac069.

31. Soreanu I, Hendler A, Dahan D et al. Marker-free genetic manipulations in yeast using CRISPR/CAS9 system. Curr Genet 2018;64:1129–39.

32. Appanah R, Lones EC, Aiello U et al. Sen1 Is Recruited to Replication Forks via Ctf4 and Mrc1 and Promotes Genome Stability. Cell Rep 2020;30:2094–2105.e9.

33. Felipe-Abrio I, Lafuente-Barquero J, García-Rubio ML et al. RNA polymerase II contributes to preventing transcription-mediated replication fork stalls. EMBO J 2015;34:236–50.

34. Weiβ M, Chanou A, Schauer T et al. Single-copy locus proteomics of early- and late-firing DNA replication origins identifies a role of Ask1/DASH complex in replication timing control. Cell Rep 2023;42:112045.

35. Cankorur-Cetinkaya A, Dereli E, Eraslan S et al. A novel strategy for selection and validation of reference genes in dynamic multidimensional experimental design in yeast. PLoS One 2012;7:e38351.

36. Dovrat D, Dahan D, Sherman S et al. A Live-Cell Imaging Approach for Measuring DNA Replication Rates. Cell Rep 2018;24:252–8.

37. Ginno PA, Lott PL, Christensen HC et al. R-Loop Formation Is a Distinctive Characteristic of Unmethylated Human CpG Island Promoters. Mol Cell 2012;45:814–25.

38. Brüning J-G, Marians KJ. Replisome bypass of transcription complexes and R-loops. Nucleic Acids Res 2020;48:10353–67.

39. Hodson C, van Twest S, Dylewska M et al. Branchpoint translocation by fork remodelers as a general mechanism of R-loop removal. Cell Rep 2022;41:111749.

40. Kumar C, Remus D. A transcription-based approach to purify R-loop-containing plasmid DNA templates in vitro. STAR Protoc 2023;4:101937.

41. Stoy H, Zwicky K, Kuster D et al. Direct visualization of transcription-replication conflicts reveals post-replicative DNA:RNA hybrids. Nat Struct Mol Biol 2023;30:348–59.

42. Laspata N, Kaur P, Mersaoui SY et al. PARP1 associates with R-loops to promote their resolution and genome stability. Nucleic Acids Res 2023;51:2215–37.

43. Kumar C, Batra S, Griffith JD et al. The interplay of RNA:DNA hybrid structure and G-quadruplexes determines the outcome of R-loop-replisome collisions. Stillman B, Tyler JK, Stillman B, et al. (eds.). Elife 2021;10:e72286.

44. Werner M, Trauner M, Schauer T et al. Transcription-replication conflicts drive R-loop-dependent nucleosome eviction and require DOT1L activity for transcription recovery. Nucleic Acids Res 2025;53, DOI: 10.1093/nar/gkaf109.

45. Lenstra TL, Coulon A, Chow CC et al. Single-Molecule Imaging Reveals a Switch between Spurious and Functional ncRNA Transcription. Mol Cell 2015;60:597–610.

46. Bruno F, Coronel-Guisado C, González-Aguilera C. Collisions of RNA polymerases behind the replication fork promote alternative RNA splicing in newly replicated chromatin. Mol Cell 2024;84:221–233.e6.

47. Voichek Y, Bar-Ziv R, Barkai N. Expression homeostasis during DNA replication. Science 2016;351:1087–90.

48. Rodríguez-Gil A, García-Martínez J, Pelechano V et al. The distribution of active RNA polymerase II along the transcribed region is gene-specific and controlled by elongation factors. Nucleic Acids Res 2010;38:4651–64.

49. Arudchandran A, Cerritelli S, Narimatsu S et al. The absence of ribonuclease H1 or H2 alters the sensitivity of Saccharomyces cerevisiae to hydroxyurea, caffeine and ethyl methanesulphonate: implications for roles of RNases H in DNA replication and repair. Genes Cells 2000;5:789–802.

50. Skourti-Stathaki K, Proudfoot NJ, Gromak N. Human senataxin resolves RNA/DNA hybrids formed at transcriptional pause sites to promote Xrn2-dependent termination. Mol Cell 2011;42:794–805.

51. Mischo HE, Gómez-González B, Grzechnik P et al. Yeast Sen1 helicase protects the genome from transcription-associated instability. Mol Cell 2011;41:21–32.

52. Alzu A, Bermejo R, Begnis M et al. Senataxin associates with replication forks to protect fork integrity across RNA-polymerase-II-transcribed genes. Cell 2012;151:835–46.

53. Schürer KA, Rudolph C, Ulrich HD et al. Yeast MPH1 gene functions in an error-free DNA damage bypass pathway that requires genes from Homologous recombination, but not from postreplicative repair. Genetics 2004;166:1673–86.

54. Lafuente-Barquero J, Luke-Glaser S, Graf M et al. The Smc5/6 complex regulates the yeast Mph1 helicase at RNA-DNA hybrid-mediated DNA damage. PLoS Genet 2017;13:e1007136.

55. Scheller J, Schürer A, Rudolph C et al. MPH1, a yeast gene encoding a DEAH protein, plays a role in protection of the genome from spontaneous and chemically induced damage. Genetics 2000;155:1069–81.

56. Kumar C, Remus D. Looping out of control: R-loops in transcription-replication conflict. Chromosoma 2024;133:37–56.

57. Prado F, Aguilera A. Impairment of replication fork progression mediates RNA polII transcription-associated recombination. EMBO J 2005;24:1267–76.

58. Wellinger RE, Prado F, Aguilera A. Replication fork progression is impaired by transcription in hyperrecombinant yeast cells lacking a functional THO complex. Mol Cell Biol 2006;26:3327–34.

59. García-Rubio M, Aguilera P, Lafuente-Barquero J et al. Yra1-bound RNA-DNA hybrids cause orientation-independent transcription-replication collisions and telomere instability. Genes Dev 2018;32:965–77.

60. Belotserkovskii BP, Shin JHS, Hanawalt PC. Strong transcription blockage mediated by R-loop formation within a G-rich homopurine-homopyrimidine sequence localized in the vicinity of the promoter. Nucleic Acids Res 2017;45:6589–99.

61. Herrera-Moyano E, Mergui X, García-Rubio ML et al. The yeast and human FACT chromatin-reorganizing complexes solve R-loop-mediated transcription-replication conflicts. Genes Dev 2014;28:735–48.

62. Lang KS, Merrikh H. Topological stress is responsible for the detrimental outcomes of head-on replication-transcription conflicts. Cell Rep 2021;34:108797.

63. Belotserkovskii BP, Tornaletti S, D’Souza AD et al. R-loop generation during transcription: Formation, processing and cellular outcomes. DNA Repair (Amst) 2018;71:69–81.

64. Kireeva ML, Komissarova N, Kashlev M. Overextended RNA:DNA hybrid as a negative regulator of RNA polymerase II processivity. J Mol Biol 2000;299:325–35.

65. Chakraborty P, Huang JTJ, Hiom K. DHX9 helicase promotes R-loop formation in cells with impaired RNA splicing. Nat Commun 2018;9:4346.

66. El Hage A, French SL, Beyer AL et al. Loss of Topoisomerase I leads to R-loop-mediated transcriptional blocks during ribosomal RNA synthesis. Genes Dev 2010;24:1546–58.

67. Bubeck D, Reijns MAM, Graham SC et al. PCNA directs type 2 RNase H activity on DNA replication and repair substrates. Nucleic Acids Res 2011;39:3652–66.

68. Nguyen HD, Yadav T, Giri S et al. Functions of Replication Protein A as a Sensor of R Loops and a Regulator of RNaseH1. Mol Cell 2017;65:832–847.e4.

69. Bayona-Feliu A, Barroso S, Muñoz S et al. The SWI/SNF chromatin remodeling complex helps resolve R-loop-mediated transcription-replication conflicts. Nat Genet 2021, DOI: 10.1038/s41588-021-00867-2.

70. Prendergast L, McClurg UL, Hristova R et al. Resolution of R-loops by INO80 promotes DNA replication and maintains cancer cell proliferation and viability. Nat Commun 2020;11:4534.

71. Lynskey ML, Brown EE, Bhargava R et al. HIRA protects telomeres against R-loop-induced instability in ALT cancer cells. Cell Rep 2024;43:114964.

72. Shin J-H, Kelman Z. The replicative helicases of bacteria, archaea, and eukarya can unwind RNA-DNA hybrid substrates. J Biol Chem 2006;281:26914–21.

73. El Hage A, Webb S, Kerr A et al. Genome-wide distribution of RNA-DNA hybrids identifies RNase H targets in tRNA genes, retrotransposons and mitochondria. PLoS Genet 2014;10:e1004716.

74. Hidmi O, Oster S, Monin J et al. TOP1 and R-loops facilitate transcriptional DSBs at hypertranscribed cancer driver genes. iScience 2024;27:109082.

