## Supplemental Information for "Replisome progression regulates R-loop mediated transcriptional repression"

### SUPPLEMENTARY INFORMATION

#### Supplementary Figures

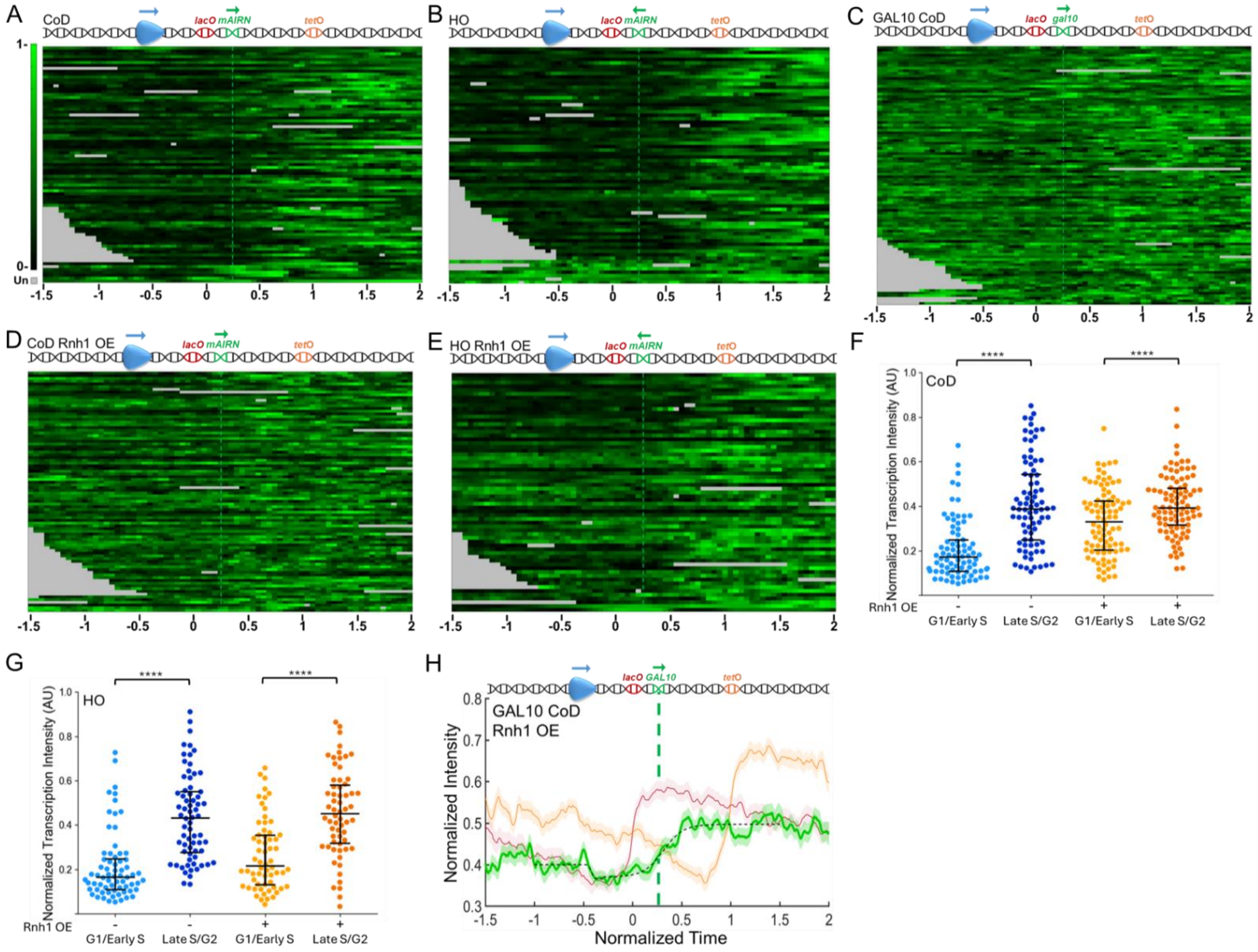


**Figure S1. Live-cell imaging of mAIRN nascent transcripts in single *Saccharomyces cerevisiae* cells.** (**A-E**) Kymographs depicting PCP-Envy foci intensity through time in (**A**) CoD, (**B**) HO, (**C**) GAL10 CoD, (**D**) CoD Rnh1 OE, and (**E**) HO Rnh1 OE yeast strains. The colour of each bin represents the strength of the PCP-Envy foci normalized fluorescent intensity in single cells (horizontal lines), with the greener the bin colour, the stronger the intensity. The time axis is normalized according to the duplication times of the arrays. Grey bins indicate timepoints that the PCP-Envy foci could not be identified (Un). Top: Scheme showing the relative positions of *ARS413*, *lacO and tetO* arrays, and mAIRN or GAL10 reporters relative to replisome movement on this locus. (**F-G**) Comparison of average single-cell PCP-Envy foci intensities in G1/Early S (before mAIRN replication) or Late S/G2 (post mAIRN replication) phases in WT or Rnh1 OE cells of (**F**) CoD or (**G**) HO yeast strains. (**H**) Same analysis as in **Fig. 2**, but for a yeast strain integrated with the Pgal-14xPP7-GAL10 cassette between the *lacO and tetO* arrays in CoD orientation with Rnh1 OE. Median replication time and cell number are listed in **Table S6**. See also [1]. **** = $p < 0.001$, n/s = not significant

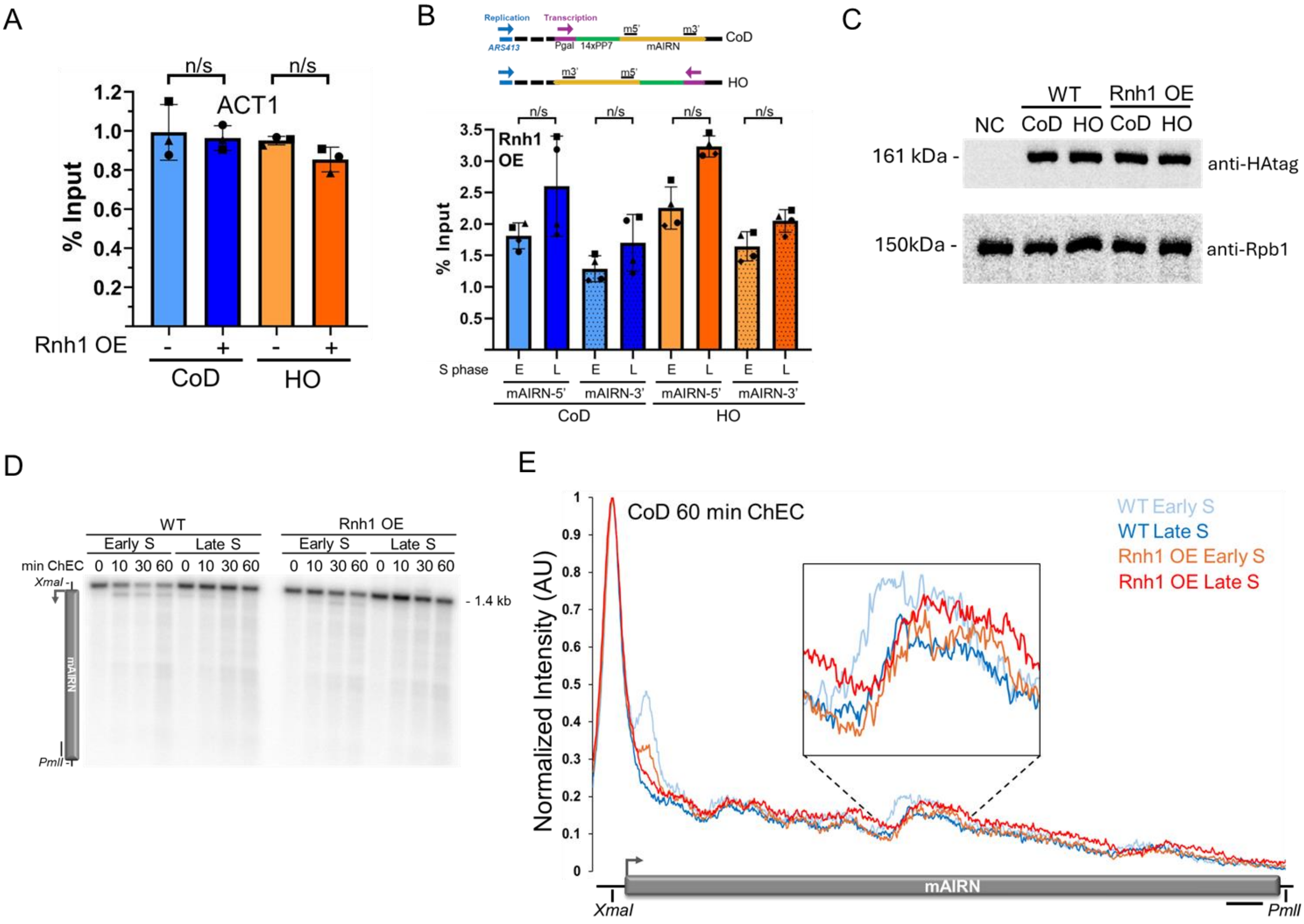


**Figure S2. Analysis of RNAPII occupancy on mAIRN.** (**A**) RNAPII ChIP-qPCR analysis, as in **Fig. 3A**, shows no RNAPII occupancy changes on ACT1 gene between WT and Rnh1 OE cells. Averages of three biological replicates are shown. Error bars represent ±SD. (**B**) Same as **Fig. 3C**, but for CoD and HO Rnh1 OE yeast strains. Averages of 4 biological replicates are shown. Error bars represent ±SD. (**C**) WB analysis of CoD and HO, WT or Rnh1 OE cells expressing Rpb2 fused with MNase and labeled with 3xHA compared to untagged control cells (NC). Anti-Rpb1 was used as a loading control. (**D-E**) Same as in **Fig. 3D&E,** for yeast strains expressing mAIRN in CoD orientation. One of two biological replicates is shown.

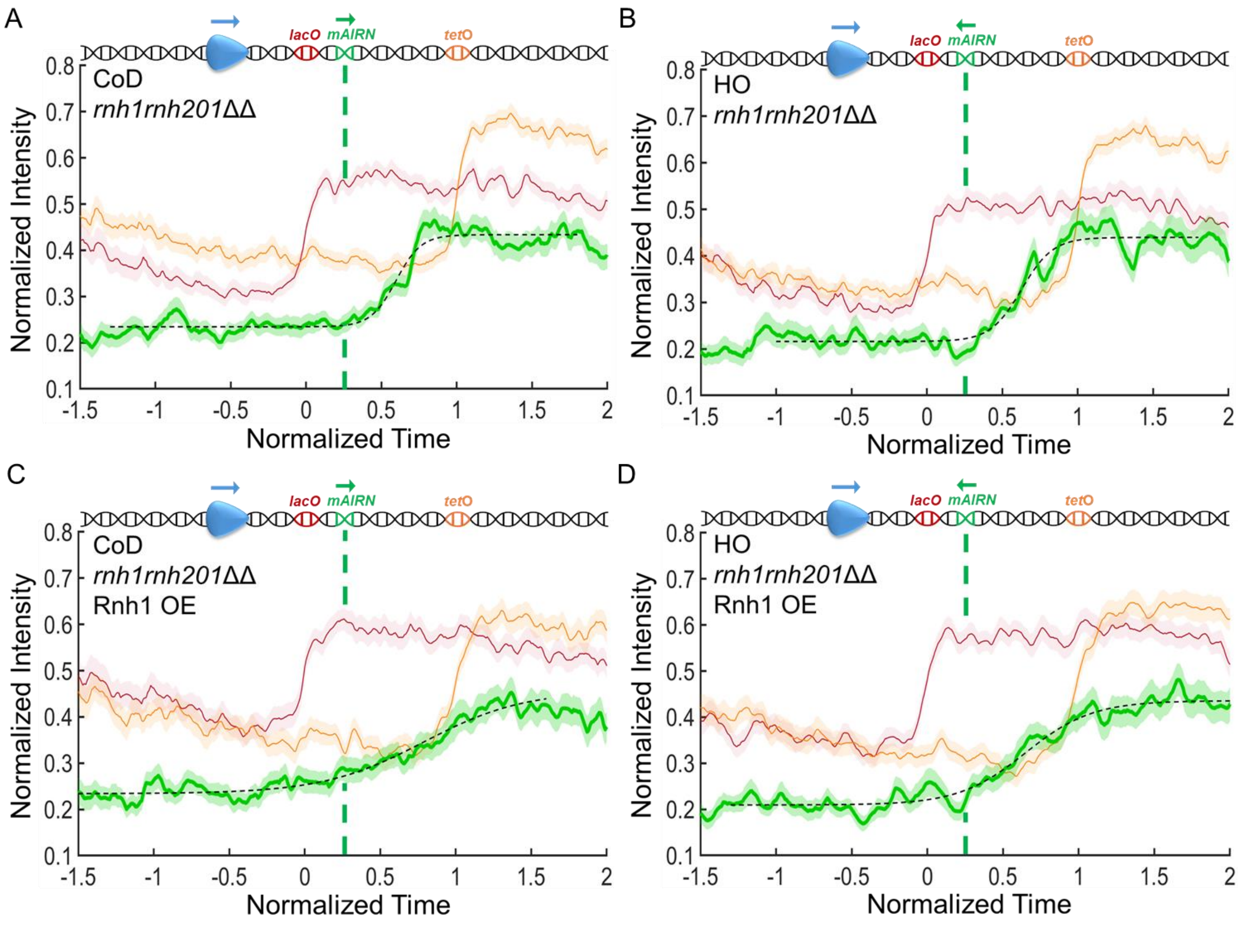


**Figure S3. mAIRN transcription profile of *rnh1rnh201*ΔΔ yeast cells during cell cycle.** (**A-D**) Average normalized fluorescent intensity of the transcription site (PCP-Envy focus, green) and the labelled *lacO* and *tetO* arrays (lacI-Halo-SiR and tetR-tdTomaro foci, red and orange respectively) were plotted over time for cells transcribing mAIRN (Induced) in (**A** and **C**) CoD or (**B** and **D**) HO orientation in *rnh1rnh201*ΔΔ yeast cells without (**A** and **B**) or with (**C** and **D**) Rnh1 OE. The time axis is normalized according to the duplication times of the arrays. Shaded areas represent SEM. A green dashed vertical line represents mAIRN duplication time. Dashed black lines represent a fit of the transcription intensity data to a sigmoidal function (see **Table S7** for parameters). Top: Scheme showing the relative positions of *ARS413*, *lacO* and *tetO* arrays, and mAIRN relative to replisome movement on this locus.

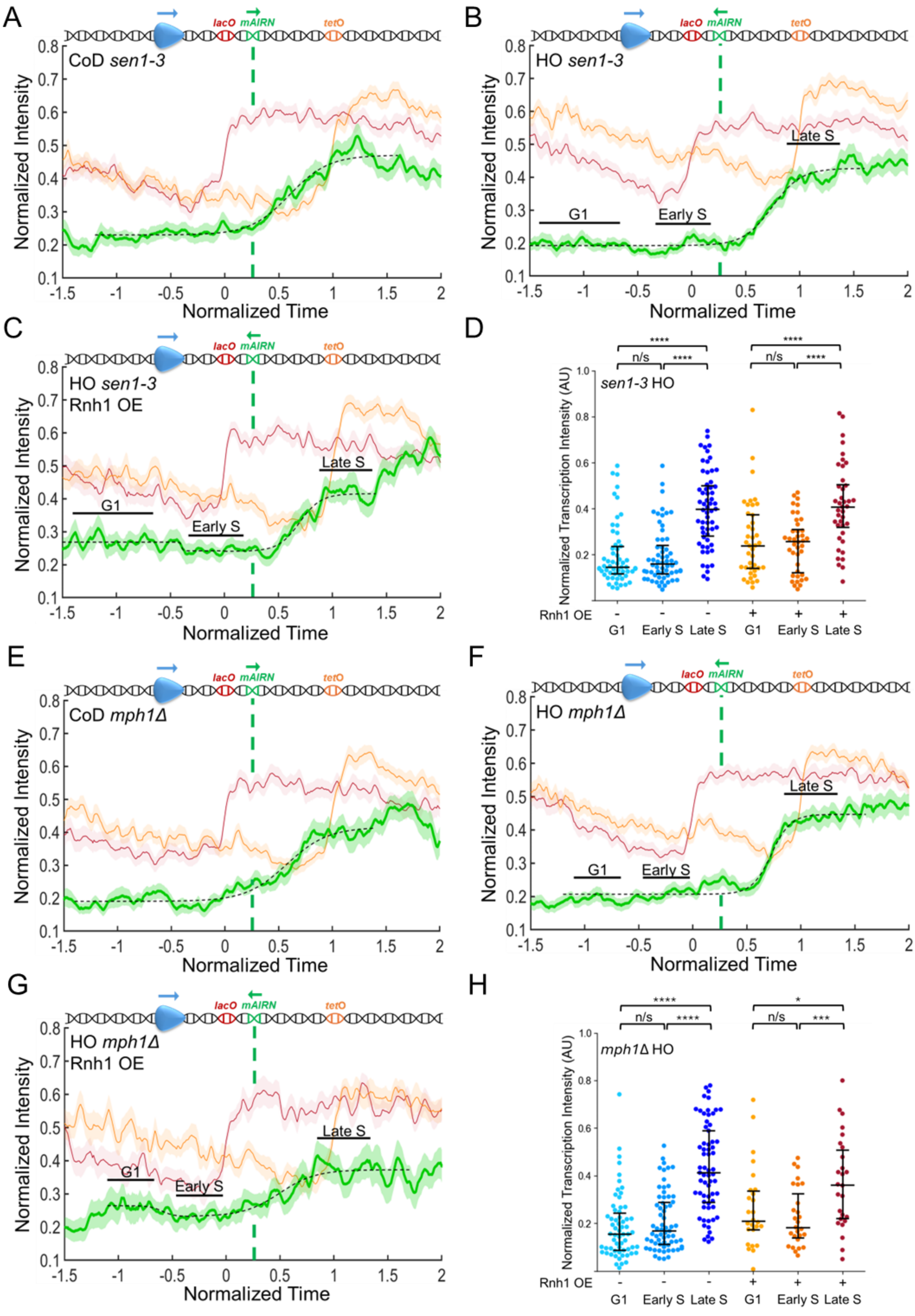
A
lacO
mAIRN
tetO
CoD sen1-3
Normalized Intensity
Normalized Time
B
HO sen1-3
G1
Early S
Late S
C
HO sen1-3
Rnh1 OE
D
sen1-3 HO
Normalized Transcription Intensity (AU)
n/s
****
Rnh1 OE
G1
Early S
Late S
E
CoD mph1Δ
F
HO mph1Δ
G
HO mph1Δ
Rnh1 OE
H
mph1Δ HO
*
***

**Figure S4. mAIRN transcription analysis of *sen1-3* and *mph1*Δ *yeast cells* during cell cycle*.*** (**A-C**) Average normalized fluorescent intensity of the transcription site (PCP-Envy focus, green) and the labelled *lacO* and *tetO* arrays (lacI-Halo-SiR and tetR-tdTomaro foci, red and orange respectively) were plotted over time for cells transcribing mAIRN (Induced) in (**A**) CoD or (**B** and **C**) HO orientation *sen1-3* yeast cells without (**A** and **B**) or with (**C**) Rnh1 OE. The time axis is normalized according to the duplication times of the arrays. Shaded areas represent SEM. A green dashed vertical line represents mAIRN duplication time. Dashed black lines represent a fit of the transcription intensity data to a sigmoidal function (see **Table S7** for parameters). Top: Scheme showing the relative positions of *ARS413*, *lacO* and *tetO* arrays, and mAIRN relative to replisome movement on this locus. (**D**) Comparison of average single-cell PCP-Envy foci intensities of WT or Rnh1 OE *sen1-3* HO yeast in G1, Early S, and Late S phase**.** (**E-H**) Same as in **A-D**, but for *mph1*Δ yeast cells, **** = p < 0.001, n/s = not significant.

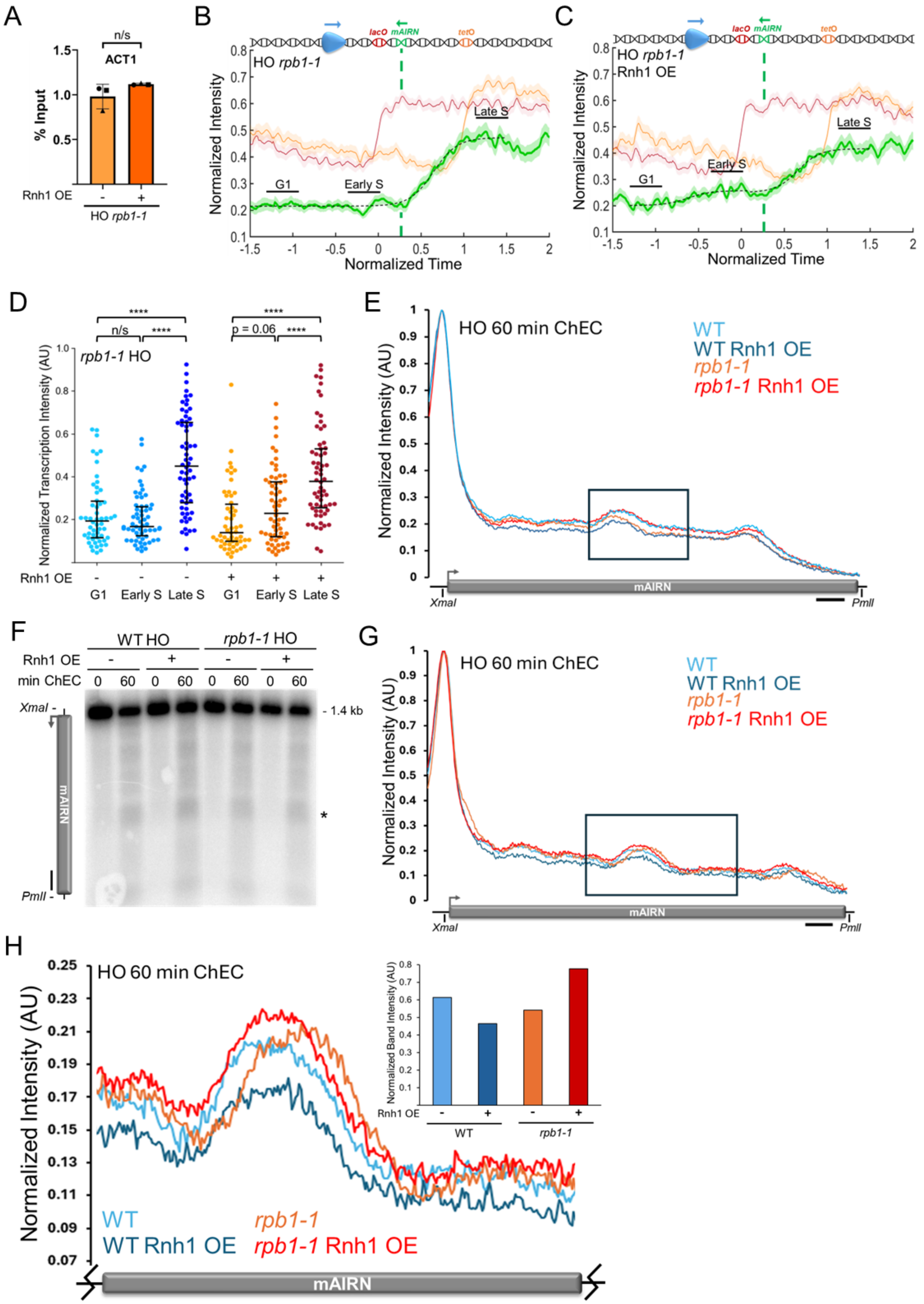
A
n/s
ACT1
% Input
Rnh1 OE
HO rpb1-1
B
lacO mAIRN tetO
HO rpb1-1
Normalized Intensity
G1
Early S
Late S
Normalized Time
C
HO rpb1-1
Rnh1 OE
Normalized Intensity
G1
Early S
Late S
Normalized Time
D
rpb1-1 HO
Normalized Transcription Intensity (AU)
n/s
p = 0.06
Rnh1 OE
G1 Early S Late S G1 Early S Late S
E
HO 60 min ChEC
WT
WT Rnh1 OE
rpb1-1
rpb1-1 Rnh1 OE
Normalized Intensity (AU)
mAIRN
XmaI
PmlI
F
WT HO
rpb1-1 HO
Rnh1 OE
min ChEC
- 1.4 kb
XmaI
mAIRN
PmlI
G
HO 60 min ChEC
WT
WT Rnh1 OE
rpb1-1
rpb1-1 Rnh1 OE
Normalized Intensity (AU)
mAIRN
XmaI
PmlI
H
HO 60 min ChEC
Normalized Intensity (AU)
Normalized Band Intensity (AU)
Rnh1 OE
WT
rpb1-1
WT
WT Rnh1 OE
rpb1-1
rpb1-1 Rnh1 OE
mAIRN

**Figure S5. mAIRN transcription analysis and RNAPII occupancy profile of *rpb1-1 yeast cells*.** (**A**) RNAPII ChIP-qPCR analysis, as in **Fig. 5A**, shows no RNAPII occupancy changes on ACT1 gene between WT and Rnh1 OE *rpb1-1* HO cells. Averages of three biological replicates are shown. Error bars represent ±SD. (**B-C**) Average normalized fluorescent intensity of the transcription site (PCP-Envy focus, green) and the labelled *lacO* and *tetO* arrays (lacI-Halo-SiR and tetR-tdTomaro foci, red and orange respectively) were plotted over time for cells transcribing mAIRN (Induced) in HO *rpb1-1* yeast cells without (**B**) or with (**C**) Rnh1 OE. The time axis is normalized according to the duplication times of the arrays. Shaded areas represent SEM. A green dashed vertical line represents mAIRN duplication time. Dashed black lines represent a fit of the transcription intensity data to a sigmoidal function (see **Table S7** for parameters). Top: Scheme showing the relative positions of *ARS413*, *lacO* and *tetO* arrays, and mAIRN relative to replisome movement on this locus. (**D**) Comparison of average single-cell PCP-Envy foci intensities of WT or Rnh1 OE *rpb1-1* HO yeast cells in G1, Early S, and Late S phase. (**E**) Relative RNAPII occupancy on the mAIRN region in HO WT or *rpb1-1* yeast cells as quantified from **Fig. 5C**. Magnification of the highlighted region is shown in **Fig. 5D**. Bottom: Scheme of mAIRN sequence aligned with the relative RNAPII occupancy quantification plot. XmaI and PmlI restriction sites and mAIRN probe relative positions are also depicted. (**F-H**) Same as in **Fig. 5C&D** and **Fig. S5E**, but for the second biological replicate, also showing elevated RNAPII occupancy after Rnh1 OE in the HO mAIRN *rpb1-1* yeast strain.

**Table S1: Yeast and bacterial strains used in this study**

| Identifier | Strain | Genotype | Source |
|---|---|---|---|
| Y0152 | mAIRN CoD | *Saccharomyces cerevisiae* W303; MATa; leu2-3,112 trp1-1; can1-100; ura3-1; ade2-1; *chrIV*:332961::*lacO*x256-TRP; *chrIV*:352558::*tetO*x224-LEU; *ade1*Δ::tetR-tdTomato+PCP-Envy; *pdr5*Δ::lacI-HaloTag; *chrIV:*336186::CUT60ter-Pgal-PP7x14-mAIRN-ADH1ter transcribed in **CoD** relative to the replisome moving from ARS413 | This study |
| Y0153 | mAIRN HO | Same as Y0001, with *chrIV*:336186::CUT60ter-Pgal-PP7x14-mAIRN-ADH1ter transcribed in **HO** relative to the replisome moving from ARS413 | This study |
| Y0167 | mAIRN CoD Rnh1 OE | Same as Y0001, with *ura3-1*::URA3-Pgal-RNH1-3HA-CYC1ter | This study |
| Y0168 | mAIRN HO Rnh1 OE | Same as Y0002, with *ura3-1*::URA3-Pgal-RNH1-3HA-CYC1ter | This study |
| Y0169 | GAL10orf CoD Rnh1 OE | Same as Y0001, with *chrIV*:336186::CUT60ter-Pgal-PP7x14-GAL10orf-CUT60ter transcribed in **CoD** relative to the replisome moving from ARS413; *ura3-1*::URA3-Pgal-RNH1-3HA-CYC1ter | This study |
| Y0156 | mAIRN CoD *rnh1rnh201*ΔΔ | Same as Y0001, but with *rnh1*Δ::*natMX*; *rnh201*Δ::*hphMX* | This study |
| Y0157 | mAIRN HO *rnh1rnh201*ΔΔ | Same as Y0002, but with *rnh1*Δ::*natMX*; *rnh201*Δ::*hphMX* | This study |
| Y0163 | mAIRN CoD *rnh1rnh201*ΔΔ Rnh1 OE | Same as Y0006, but with *natMX*::Pgal-RNH1-3HA-CYC1ter | This study |
| Y0164 | mAIRN HO *rnh1rnh201*ΔΔ Rnh1 OE | Same as Y0007, but with *natMX*::Pgal-RNH1-3HA-CYC1ter | This study |
| Y0162 | mAIRN CoD *sen1-3* | Same as Y0001, with SEN1::*sen1-3 (W773A, E774A, W777A)* | This study |
| Y0161 | mAIRN HO *sen1-3* | Same as Y0002, with SEN1::*sen1-3 (W773A, E774A, W777A)* | This study |
| Y0170 | mAIRN HO *sen1-3* Rnh1 OE | Same as Y0011, with ura3-1::URA3-Pgal-RNH1-3HA-CYC1ter | This study |
| Y0165 | mAIRN CoD *mph1*Δ | Same as Y0001, with *mph1*Δ::*kanMX* | This study |
| Y0166 | mAIRN HO *mph1*Δ | Same as Y0002, with *mph1*Δ::*kanMX* | This study |
| Y0173 | mAIRN HO *mph1*Δ Rnh1 OE | Same as Y0014, with ura3-1::URA3-Pgal-RNH1-3HA-CYC1ter | This study |
| Y0175 | mAIRN HO *rpb1-1* | Same as Y002, with RPB1::*rpb1-1 (G4622A)* | This study |
| Y0177 | mAIRN HO *rpb1-1* Rnh1 OE | Same as Y0016, with ura3-1::URA3-Pgal-RNH1-3HA-CYC1ter | This study |
| B0001 (Cat# 18265017) | *Escherichia coli* DH5alpha | - | Thermo Fischer Scientific |

**Table S2: Plasmids used in this study**

| Identifier | Plasmid | Source |
|---|---|---|
| K349 | pAG25-C60ter-Pgal-14xPP7-mAIRN-ADH1ter | This study |
| K346 | pTL-Pgal-RNH1-3HA-CYC1ter | This study |
| K350 | pCAS9-RPB1-gRNA2 | Tsirkas et al. [1] |
| K351 | pCAS9-SEN1-gRNA1 | This study |
| K142 | MNase-3xHA | This study |

**Table S3: Oligos used in the study.**

| Name | Sequence (5'-3') |
|---|---|
| mAIRN5' Frw | tagaggattccgcaaaggaa |
| mAIRN5' Rev | ttcaccctagcgctgaatct |
| mAIRN3' Frw | cgagagaggctaagggtgaa |
| mAIRN3' Rev | acatggtcctgctggagttc |
| ACT1orf Frw | agagttgccccagaagaaca |
| ACT1orf Rev | ggcttggatggaaacgtaga |
| TDH3orf Frw | caaggaaaccacctacga |
| TDH3orf Rev | cgaagatggaagagtgagag |
| Spike-in Frw (M13 Frw) | gtaaaacgacggccagt |
| Spike-in Rev (M13 Rev) | caggaaacagctatgac |
| mAIRN-3-SB-Frw | gctcagagggttccgagctatcc |
| mAIRN-3-SB-Rev | gttcatctctcctgtaacatggcac |

**Table S4: Reagents and commercial kits/assays used in this study**

| Reagent/Material | Source | Identifier |
|---|---|---|
| Anti-HA-Peroxidase, high affinity (3F10) | Roche | Cat# 12013819001; RRID:AB_390917 |
| Mouse monoclonal anti-RNA Polymerase II CTD Antibody (8WG16) | Millipore | Cat# 05-952; RRID:AB_11213782 |
| Polyclonal RNA polymerase II CTD repeat YSPTSPS (phospho S2) | Abcam | Cat# ab193468; RRID:AB_290555 |
| Goat Anti-Rabbit IgG (H+L), Horseradish peroxidase conjugate | Invitrogen | Cat# G21234; RRID:AB_2536530 |
| Yeast Alpha-Factor Mating Pheromone | Genscript | Cat# RP01002 |
| Concanavalin A | Sigma-Aldrich | Cat# L7647 |
| Silicon Rhodamine-HALO dye | Tsirkas et al. [1] | N/A |
| Polyethylene Glycol 3350 | Sigma-Aldrich | Cat# P4338 |
| Lithium acetate dihydrate | Sigma-Aldrich | Cat# L6883 |
| Herring Sperm DNA | Promega | Cat# D1816 |
| Lithium Chloride | CarlRoth | Cat# 2312123 |
| EGTA | SantaCruz | Cat# Sc-3593A |
| Tris Base | ChemCruz | Cat# sc-3715B |
| Formaldehyde | Thermo Fisher Scientific | Cat# 28908 |
| Pierce Protein A Magnetic Beads | Thermo Fisher Scientific | Cat# 10001D |
| Deoxycholic acid | Santa Cruz | Cat# sc-214865A |
| Triton X-100 | Sigma-Aldrich | Cat# X100-100ML |
| Spermidine | Sigma-Aldrich | Cat# S0266-1G |
| Spermine | Sigma-Aldrich | Cat# S3256 |
| SYBR Safe DNA Gel Stain | Invitrogen | Cat# S33102 |

| | | |
|---|---|---|
| Ammonium Acetate | SERVA | Cat# 39750.01 |
| Hydrochloric Acid 5N | AppliChem | Cat# 182109.1211 |
| Sodium Hydroxide | Sigma-Aldrich | Cat# S5881 |
| Sodium Chloride | Supelco | Cat# 1.06404.5000 |
| DNase I | New England Biolabs | Cat# M0303S |
| RNase A | Thermo Fisher Scientific | Cat# EN0531 |
| Proteinase K | SERVA | Cat# 33756 |
| Phenol:chlorophorm:isoamyl alcohol (25:24:1; v/v) | Invitrogen | Cat# 15593-031 |
| OmniPur Phenol:Chloroform:Isoamyl Alcohol, 25:24:1 | Millipore | Cat# 6805-100ML |
| Glycogen | Thermo Fisher Scientific | Cat# AM9510 |
| Igepal CA-630 | Sigma-Aldrich | Cat# I3021-100ML |
| Halt Protease and Phosphatase Inhibitor Cocktail (100x) | Thermo Fisher Scientific | Cat# 78446 |
| BSA Fraction V | Sigma-Aldrich | Cat# 10735078001 |
| Tween-20 | Kraft | Cat# 21440.2000 |
| NuPAGE 4-12% Bis-Tris Protein Gels 1.5 mm | Thermo Fisher Scientific | Cat# NP0336BOX |
| Immobilon Transfer Membrane PVDF | Millipore | Cat# IPVH00010 |
| SuperSignal™ West Pico PLUS Chemiluminescent Substrate | Thermo Fisher Scientific | Cat# 34580 |
| iTaq Universal SYBR Green Supermix | Bio-Rad | Cat# 1725121 |
| SuperScript III First-Strand Synthesis System | Thermo Fisher Scientific | Cat# 18080-051 |
| RadPrime DNA Labeling System | Invitrogen | Cat# 18428-011 |
| μ-Slide 8 Well Uncoated | Ibidi | Cat# 80821 |

**Table S5: Software and algorithms used in this study**

| Software/Algorithm | Source | Identifier |
|---|---|---|
| ImageJ | NIH | https://imagej.net/ij/index.html |
| Zen (version 3.0 blue edition) | Zeiss | https://www.micro-shop.zeiss.com/ en/us/softwarefinder/#select-soft-ware |
| Matlab (version 2019b) | MathWorks | https://www.mathworks.com/products.html?s_tid=gn_ps |
| Transcriptomatic5 | This study | 10.5281/zenodo.15601453 |
| Python (versions 2.7 and 3.8) | Python | https://www.python.org |
| PyCharm (2020.2) | JetBrains | https://www.jetbrains.com/pycharm/ |
| R v4.1.2 | R core Team | https://www.r-project.org |
| GraphPad Prism 9 | GraphPad Software | https://www.graphpad.com/ |

**Table S6: Median replication time and cell number of yeast strains for live-cell imaging experiments**

| Yeast Strain and condition | Median Replication time (min) | Cell number |
|---|---|---|
| mAIRN CoD Uninduced | 17.9 | 70 |
| mAIRN CoD Induced | 17.5 | 85 |
| mAIRN HO Uninduced | 17.4 | 69 |
| mAIRN HO Induced | 17.6 | 71 |
| mAIRN CoD Rnh1 OE Induced | 18.1 | 95 |
| mAIRN HO Rnh1 OE Induced | 17.7 | 69 |

| | | |
|---|---|---|
| GAL10orf CoD Rnh1 OE Induced | 17.0 | 96 |
| mAIRN CoD *rnh1rnh201*ΔΔ Uninduced | 16.8 | 61 |
| mAIRN CoD *rnh1rnh201*ΔΔ Induced | 18.8 | 75 |
| mAIRN HO *rnh1rnh201*ΔΔ Uninduced | 14.3 | 62 |
| mAIRN HO *rnh1rnh201*ΔΔ Induced | 17.0 | 67 |
| mAIRN CoD *rnh1rnh201*ΔΔ Rnh1 OE Induced | 17.3 | 53 |
| mAIRN HO *rnh1rnh201*ΔΔ Rnh1 OE Induced | 15.0 | 57 |
| mAIRN CoD *sen1-3* Uninduced | 16.3 | 50 |
| mAIRN CoD *sen1-3* Induced | 16.8 | 50 |
| mAIRN HO *sen1-3* Uninduced | 17.3 | 59 |
| mAIRN HO *sen1-3* Induced | 19.4 | 61 |
| mAIRN HO *sen1-3* Rnh1 OE Induced | 16.7 | 55 |
| mAIRN CoD *mph1*Δ Induced | 17.1 | 50 |
| mAIRN HO *mph1*Δ Induced | 20.2 | 66 |
| mAIRN HO *mph1*Δ Rnh1 OE Induced | 15.7 | 32 |
| mAIRN HO *rpb1-1* Induced | 17.1 | 70 |
| mAIRN HO *rpb1-1* Rnh1 OE Induced | 20.1 | 66 |

**Table S7: Distance of PCP-Envy foci intensity transcription increase upon mAIRN replication and the calculated slope of the sigmoid plot fitted on this transcriptional increase.**

| **Yeast Strain/Condition** | **Distance (kb) of mAIRN transcription increase following mAIRN replication**[a] | **Sigmoid Increase Slope**[b] |
|---|---|---|
| mAIRN CoD | 9.3 | 4.5 |
| mAIRN HO | 17.7 | 7.2 |
| mAIRN CoD rnh1rnh201ΔΔ | 10.00 | 12.0 |
| mAIRN CoD rnh1rnh201ΔΔ Rnh1 OE | 16.30 | 3.0 |

| | | |
|---|---|---|
| mAIRN HO rnh1rnh201ΔΔ | 11.00 | 9.6 |
| mAIRN HO rnh1rnh201ΔΔ Rnh1 OE | 11.60 | 4.5 |

[a] Calculated as the distance between the middle of PP7-mAIRN reporter cassette and the midpoint of the sigmoid function plotted on the PCP-Envy intensity increase

[b] A high slope indicates a delayed/longer increase from the minimum to the maximum of the sigmoid plot

#### REFERENCES

1. Tsirkas I, Dovrat D, Thangaraj M et al. Transcription-replication coordination revealed in single live cells. Nucleic Acids Res 2022, DOI: 10.1093/nar/gkac069.
